# Methods for Copy Number Aberration Detection from Single-cell DNA Sequencing Data

**DOI:** 10.1101/696179

**Authors:** Xian Fan, Mohammadamin Edrisi, Nicholas Navin, Luay Nakhleh

**Author notes:** Contributed equally (Xian Fan, Mohammadamin Edrisi).

## Abstract

Single-cell DNA sequencing technologies are enabling the study of mutations and their evolutionary trajectories in cancer. Somatic copy number aberrations (CNAs) have been implicated in the development and progression of various types of cancer. A wide array of methods for CNA detection has been either developed specifically for or adapted to single-cell DNA sequencing data. Understanding the strengths and limitations that are unique to each of these methods is very important for obtaining accurate copy number profiles from single-cell DNA sequencing data. Here we review the major steps that are followed by these methods when analyzing such data, and then review the strengths and limitations of the methods individually. In terms of segmenting the genome into regions of different copy numbers, we categorize the methods into three groups, select a representative method from each group that has been commonly used in this context, and benchmark them on simulated as well as real datasets. While single-cell DNA sequencing is very promising for elucidating and understanding CNAs, even the best existing method does not exceed 80% accuracy. New methods that significantly improve upon the accuracy of these three methods are needed. Furthermore, with the large datasets being generated, the methods must be computationally efficient.

---

Acquired mutations are the main causes of cancer [19, 42, 62]. Copy number aberrations (CNAs) are one such type of acquired mutations and have been implicated in over-regulating oncogenes or down-regulating tumor suppressor genes [9]. Consequently, accurate detection of CNAs could hold a great promise to understanding some of the genetic underpinnings of cancer as well as developing targeted drugs. In the past two decades, a wide array of DNA technologies have been used to detect CNAs, among which the three most widely used are array Comparative Genomic Hybridization (aCGH), Next Generation Sequencing (NGS), and single-cell sequencing [40].

Array CGH [13] is a cytogenetic approach that uses fluorescent dyes on the test (tumor) and reference (normal) samples, and applies them to a microarray, which is an array of probes. Each probe is a DNA sequence that represents a region of interest. The size of a probe depends on the DNA sequence being used, and it varies from dozens of base pairs, such as oligonucleotides, to thousands of base pairs, such as bacterial artificial chromosomes. The probes from the paired samples, after being mixed together, hybridize at each target region. The fluorescence intensities can then be measured for both samples, and the ratio of the two is used to inform about CNAs of the test sample relative to the normal one. Array CGH data is advantageous in comprehensively detecting aneuploidies, amplifications, and deletions simultaneously. A few computational methods [27, 55, 61] have been developed to detect CNAs using aCGH data. DNAcopy [55] applies a modification of binary segmentation [60] called circular binary segmentation (CBS) to aCGH to overcome data noise, but it suffers from the problem of outliers [27, 61]. HMMcopy [61] was designed to ameliorate the problem of outliers and uses a Hidden Markov Model (HMM) to divide the genome into piecewise fixed segments in order to make inferences on CNAs. However, since aCGH data is limited in resolution and throughput [17], as well as suffers from a hybridization bias problem, it is not the optimal technology to detect CNAs for cancer samples.

Unlike aCGH, which obtains signal on only a limited number of genomic sites of interest, NGS technology makes it possible to survey the whole genome at a nucleotide-level resolution by sequencing millions of small fragments (reads) of the genome in parallel. By aligning the reads to an assembled reference genome, the reads that cover a position in the genome are counted to obtain the read depth at that position. Read depths at different regions of the genome can then be contrasted to assess hypothesized genomic locations where copy number gains and losses had occurred. NGS technologies suffer from high false positive rate compared with aCGH, due mainly to GC bias and the presence of repetitive regions [30, 50]. Even more challenging in the case of cancer genomes that are often aneuploid, contamination of normal cells may occur in the bulk tissue further complicating the task of estimating the absolute copy number from NGS data. To overcome these challenges, a plethora of computational tools [1, 11, 15, 26, 29, 45, 48–50] have been developed for detecting CNAs from NGS data. SeqCNA [50] filters out potentially false-positive CNAs and corrects GC content for a more accurate CNA detection. CNAseg [29] analyzes flowcell-to-flowcell variability to avoid false-positive CNAs. CNAnorm [26] addresses the normal contamination and aneuploidy of the tumor sample to infer CNAs accurately. Paired-end NGS data provides another modality in addition to the read depth to infer CNA, and a few bioinformatics tools use this, including, for example, CNVer [48], CNVnator [1], ReadDepth [49], and Mseq-CNV [45].

Although both aCGH and NGS data can be used to detect CNAs, they do not provide high-throughput data at the single-cell resolution that is ideal for understanding tumor growth. In particular, intratumor heterogeneity [54] is best understood by using data obtained from individual cells within the tumor tissue. Indeed, in the last ten years, the field has made great strides towards developing technologies for single-cell DNA sequencing. Data generated by these technologies can be analyzed to detect CNAs and other types of mutations in individual cells and individual clones within the heterogeneous tumor [52]. For example, DOP-PCR is a PCR amplification method that generates low-coverage data suitable for CNA detection in single-cell data [4, 6, 14, 51]. However, it also suffers from uneven coverage and allelic dropout [52] that could lead to false-positive calls of CNAs. Beyond this method, three tools have been extensively applied to single-cell sequencing data for CNA detection: AneuFinder [5, 67], CopyNumber [53], and Ginkgo [24]. Like HMMcopy, AneuFinder uses a Hidden Markov Model (HMM) to determine the segmentation of the genome and the absolute copy number of each segment. CopyNumber [53] pools all cells together for joint segmentation to improve boundary detection accuracy. Since cancer cells in the same subclone mostly share the same CNA boundaries, such a strategy can improve the nucleotide resolution of the boundary by implicitly amplifying the signal in the data. Ginkgo [24] uses Circular Binary Segmentation (CBS) [55] to segment the genome, followed by inferring the integer value of the absolute copy number. It is worth noting that some methods designed for aCGH and NGS data have also been extensively used on single-cell data [28, 34, 74, 83]. One such method is HMM-copy [36]. As both AneuFinder and HMMcopy are HMM-based methods, we focus on HMMcopy as a representative of the HMM-based approach. A more recent method is SCNV [73], which uses a bin-free segmentation method to do segmentation and copy number profiling. However, the method has not been widely applied to single-cell DNA studies. Among CopyNumber, Ginkgo, and HMMcopy, only CopyNumber utilized the pooled information from single-cell data. The other two methods can be equally well applied to bulk samples. Moreover, HMMcopy was designed for aCGH data originally, and thus does not take into account the specific error profiles that characterize single-cell sequencing data, such as low and uneven coverage, or the computational challenges that arise due to biological processes such as aneuploidy in a tumor single cell.

In this paper, we review 26 methods that have been developed for CNV and/or CNA detection. We discuss these methods in terms of several general steps that are often applied to single-cell DNA sequencing data for CNA detection, and highlight the strengths and limitations of each method, in particular with respect to their applicability to single-cell DNA data from cancer genomes. A major step in CNA detection is that of partitioning a genome into segments so that the genomic region that corresponds to a segment has a single copy number and every two adjacent segments differ in their copy numbers. There are three approaches to this step, based on a sliding window, an objective function, or HMMs. In the second part of the review, we benchmark three methods that have been widely applied to analyze single-cell data [3, 5, 23, 28, 32, 34, 41, 43, 46, 56, 57, 64, 68, 71, 74, 79, 83] and that represent the three segmentation approaches: CopyNumber, Ginkgo, and HMMcopy. For this goal, we use synthetic data generated by a specifically developed simulator of single-cell DNA data in the presence of CNAs, as well as a real dataset. In addition to assessing the accuracy of the methods in terms of precision and recall, we introduced the use of parsimony-based counting of copy number changes as a potential indicator of accuracy, especially when the ground truth is unknown. We found that in terms of accuracy of the detected breakpoints and memory consumption, HMMcopy is the best of the three methods, and in terms of running time, it is slower than CopyNumber but faster than Ginkgo. However, when evaluated the methods in terms of the actual copy number profiles they detect, we found that Ginkgo is more accurate than HMMcopy; in fact, we found that HMMcopy is not stable at this task (paradoxically, CopyNumber does not detect actual copy numbers). In terms of accuracy, CopyNumber has a very poor performance. While Knouse *et al.* [34] assessed the performance of CBS and HMM-based methods on single-cell DNA sequencing data, their evaluation is limited to CNVs in brain and skin cells. Moreover, they did not investigate the effect of the ploidy on the accuracy of the methods. Our results highlight the need for developing new accurate and efficient methods for CNA detection from single-cell DNA data. Our review of the 26 methods and highlighting their strengths and limitations can help in these further developments, as strengths of the different methods could potentially be combined to produce more powerful methods.

## General Steps of a CNA Detection Pipeline

In the absence of copy number aberrations and/or variations, a diploid genome has precisely two copies of each chromosome. However, when CNVs or CNAs occur, the number of copies varies across the genome. In this case, *segmentation* is the task of partitioning the genome into maximum-length contiguous regions such that the copy number at each site within a region is the same. Each such region is called a *segment*, and the copy number shared by all sites within that segment is referred to as the *absolute copy number*. CNA detection refers to the task of identifying the segments and absolute copy numbers across a given genome. In the case of a diploid genome with no CNVs or CNAs, there is one segment, which is the whole genome, and the absolute copy number is 2.

In general, a method for CNA detection from single-cell DNA data follows some or all of the following seven steps, illustrated in Fig. 1:

- *Binning.* Partitioning the genome into fixed- or variable-size bins so that the resolution at which the genome is segmented and copy numbers are called is defined by bins rather than by individual sites.
- *GC correction.* Correcting read counts according to the GC content in the corresponding genomic region to remove the GC content’s effect on the read counts.
- *Mappability correction.* Correcting read counts according to how mappable a genomic region is. The higher the repetitiveness in a region, the lower its mappability, and the lower the number of uniquely aligned reads.
- *Removal of outlier bins.* Identifying and excluding bins that have extremely high read count regardless of the actual copy number in such bins. These bins often occur near the centromere and telomere of each chromosome.
- *Removal of outlier cells.* Identifying and excluding cells whose read count profiles are low in signal-to-noise ratio (SNR). Such cells either have noisy read counts, or are low in sequence coverage.
- *Segmentation.* Identifying the boundaries (physical locations) between genomic regions that have different absolute copy numbers.
- *Calling the absolute copy numbers.* Inferring the actual absolute copy number within each segment.

**Figure 1:**
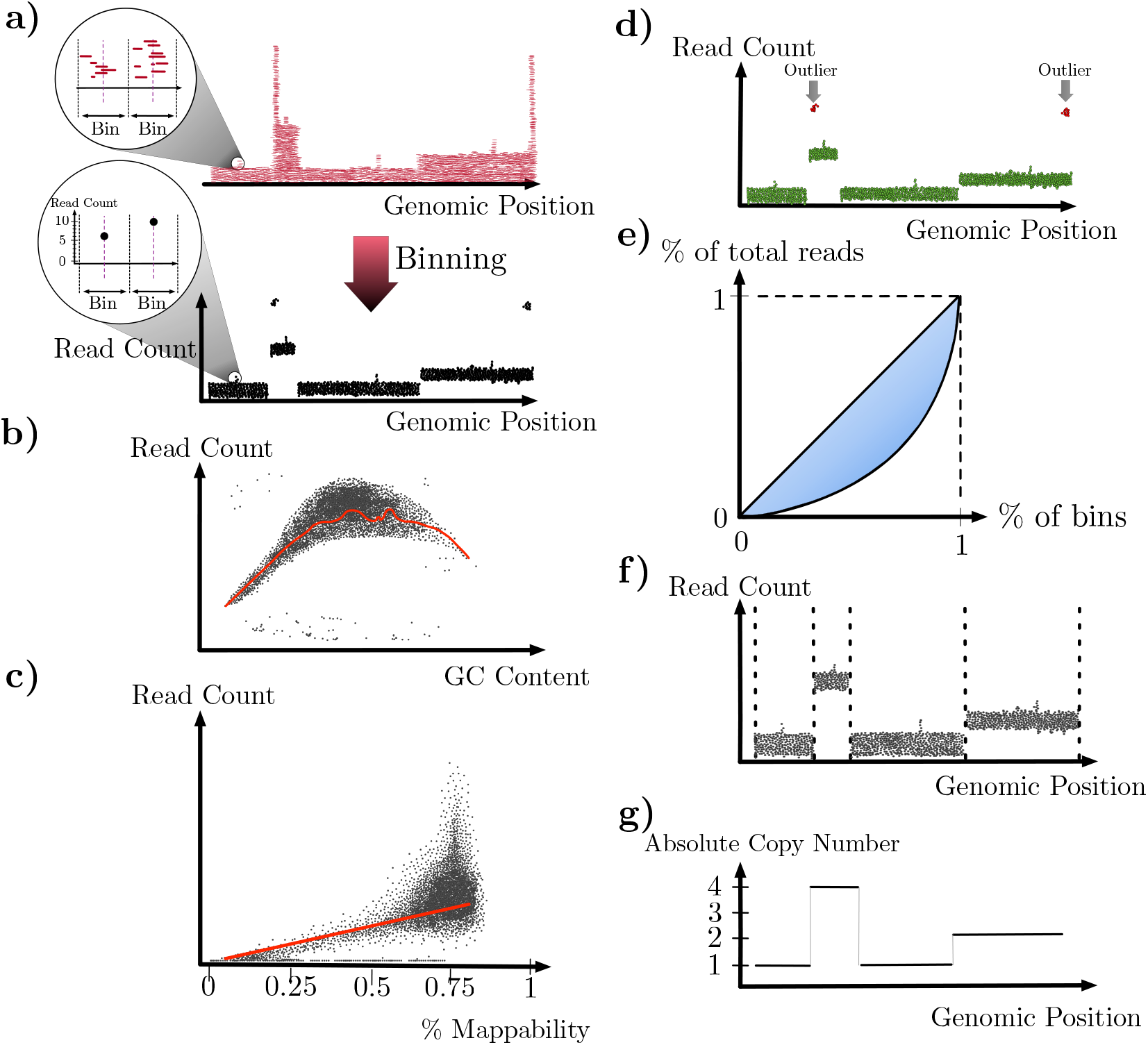
Illustration of the seven steps in CNA detection in single-cell sequencing. (a) Binning. The number of reads within each bin (bottom) is computed from the pileup of the reads according to where they align (top). (b) GC correction. Scatter plot of read count per bin with respect to the GC content of the bin. The red curve represents the corresponding regression. (c) Mappability correction. Scatter plot of read count per bin with respect to the mappability of the bin. The red curve represents the corresponding regression. (d) Removal of outlier bins. Scatter plot of read count per bin with respect to the genomic position is shown. Outlier bins are shown in red, in contrast with the rest of the genome which are in green. (e) Removal of outlier cells. A Lorenz curve for the read count at all bins is shown. Gini index is the highlighted area between the Lorenz curve and the diagonal line. The higher the Gini index, the more likely the cell is an outlier. (f) Segmentation. Scatter plot of read count per bin with respect to the genomic position is shown. Dotted vertical lines correspond to the segments’ boundaries. (g) Calling the absolute copy numbers. The copy number—a non-negative integer—for each segment is determined.

We now discuss the details of each of these seven steps, and illustrations of them in Fig. 1. We then review existing methods for CNA detection while referring to the steps they employ.

### Binning

Bins are computed so that reads within a bin are aggregated to reduce the effects of variable amplification and sequence sampling [24]. Most methods that employ binning use bins of a fixed size (all bins have the same size). However, the use of variable-size bins has been proposed [51] to avoid high false-positive deletion calls in repetitive regions, as a result of the removal of low mapping quality reads. Bins of variable sizes are determined as follows. First, the reference genome is sequenced *in silico* to produce a set of reads. The reads are then aligned back to the reference, and those that cannot be uniquely aligned are removed. The genome is then divided into variable-size bins, each bin containing roughly the same number of uniquely aligned reads. For the repetitive regions, their variable bin size is expected to be larger since the number of uniquely aligned reads is smaller [51].

While using variable-size bins effectively reduces false-positive deletion calls, it is restricted in a few aspects. First, the bin size is with respect to a specific reference genome. Different reference genomes could give rise to different binning outcomes. Second, the bin size is affected by the particular read length being simulated. More generally, determining the optimal bin size (variable or fixed) is hard due to variable read coverage in single-cell data, as mentioned in [73], since there needs to be a certain number of reads in each bin so that the resulting read counts follow a normal distribution, according to the central limit theorem. Furthermore, the use of bins could limit the segmentation resolution since bins restrict the segment boundaries to be a subset of the bin boundaries. Finally, the use of bins may generate false-negative calls for CNAs whose size is close to or smaller than the bin size as the number of the bins in that segment is so small that the signal is often mistaken as noise and will not be called as a CNA. For single-cell data, in particular, this could lead to false-negative large CNAs, as the bin size is often around 200kbp [23].

### GC correction

GC correction is a necessary step due to the dropping read coverage at the regions with extreme GC contents [8, 80], a phenomenon referred to as GC bias. As the GC content is the percentage of the nucleotides being G or C often measured within a certain genomic region, a binning step is needed to divide the whole genome into multiple bins. Thus it has been observed that a binning step always precedes a GC correction step for most of the methods. GC correction starts with modeling the change of (log normalized) read count with respect to GC content, either parametrically by a polynomial function, or non-parametrically by LOESS regression [5, 23, 24]. The data to be analyzed is used to study the model, since different sample preparation libraries may produce different curves of read count with respect to GC content. The read count at each bin is then normalized according to the model learned in the previous step to remove the effect of GC content. The output of this step is the modified read count at each bin. Most methods correct GC bias under the assumption that a majority of the genome is diploid and does not harbor copy number variations/alterations. Such an assumption, however, is not necessarily true. Specifically, tumor genomes can be aneuploid and thus contain chromosomal amplifications [23]. When no copy number dominates a genome’s copy number profile, one has to consider which copy number this bin is at when modeling its read count by the GC content [12]. To do so, a pre-segmentation and estimation of the absolute copy number is required for GC correction. However, accurate segmentation and estimation of the absolute copy number depend on accurate GC correction reciprocally. An iterative approach can be applied to update segmentation and each bin’s absolute copy number, as well as the modeling of read count with respect to the GC content [50, 72].

### Mappability correction

Mappability of a genomic region is determined by the percentage of the uniquely mappable positions in this region. Similar to GC correction, a binning step is required so that the whole genome can be divided into multiple bins for a measure of the mappability of each bin. If a bin is mainly composed of N nucleotides or repetitive regions, its mappability is possibly low. The process of determining the uniquely mappable positions is described above. Correction of the read counts according to mappability can follow the same process as GC correction, i.e., learning the curve of the read count with respect to the mappability of each bin, and normalizing the read count accordingly to remove the effect of the mappability by LOWESS regression or a parametric function. As described above, the use of variable-size bins aims to correct the uneven mappability, resulting in the same number of uniquely mappable reads in each bin.

### Removal of outlier bins

Bins that are identified as outliers are removed from further analysis by their genomic positions, GC content, or their read counts. Most often, bins at centromere or telomere of a chromosome, with extreme GC content or with zero or extremely high read counts, are identified as outliers.

### Removal of outlier cells

Cells are identified as outliers and removed from further analysis if their coverage is lower than expected [23, 73], or their Gini coefficient is larger than expected [72]. Gini coefficient measures the fluctuation of the read count across bins. The higher the Gini coefficient, the smaller the signal-to-noise ratio. Both low coverage and large Gini coefficient indicate potential failed library preparation.

### Segmentation

There are three approaches to CNA detection (Fig. 2): an HMM-based approach, a sliding-window approach, and an objective-function-based approach. In the HMM-based approach, states correspond to the different possible absolute copy numbers, and transitions between states capture the segmentation (i.e., transition out of a state at bin *i* denotes that bins *i* − 1 and *i* belong to two different segments). The sliding-window approach segments the genome by statistical testing, looking for regions whose read counts differ greatly from those of other regions. Methods that use the sliding-window approach do not calculate the absolute copy number simultaneously with segmentation. A post-processing approach is needed, which usually involves testing different candidates of ploidy and selecting the one whose resulting copy number profile is as close to integer numbers as possible. Finally, the objective-function-based approach combines the approximation to the data and limitation of breakpoints in one formula. Such approaches model the (normalized) read count by a piecewise constant function, so that the function i) is in fidelity to the data as much as possible; and ii) has as few changes as possible. To achieve this, an objective function is constructed and usually has two terms. The first term is the distance between the underlying piecewise constant function and the given data. The second term is the number of the changes, which is usually an L1 norm total variation term. The two terms are often combined by a Lagrange multiplier. Numerical methods are often applied to find the piecewise constant function that minimizes the objective function. Methods based on the objective-function approach do not simultaneously assign absolute copy number to each segment; a post-processing step is needed to assign breakpoints between segments as well as the absolute copy number at each segment.

**Figure 2:**
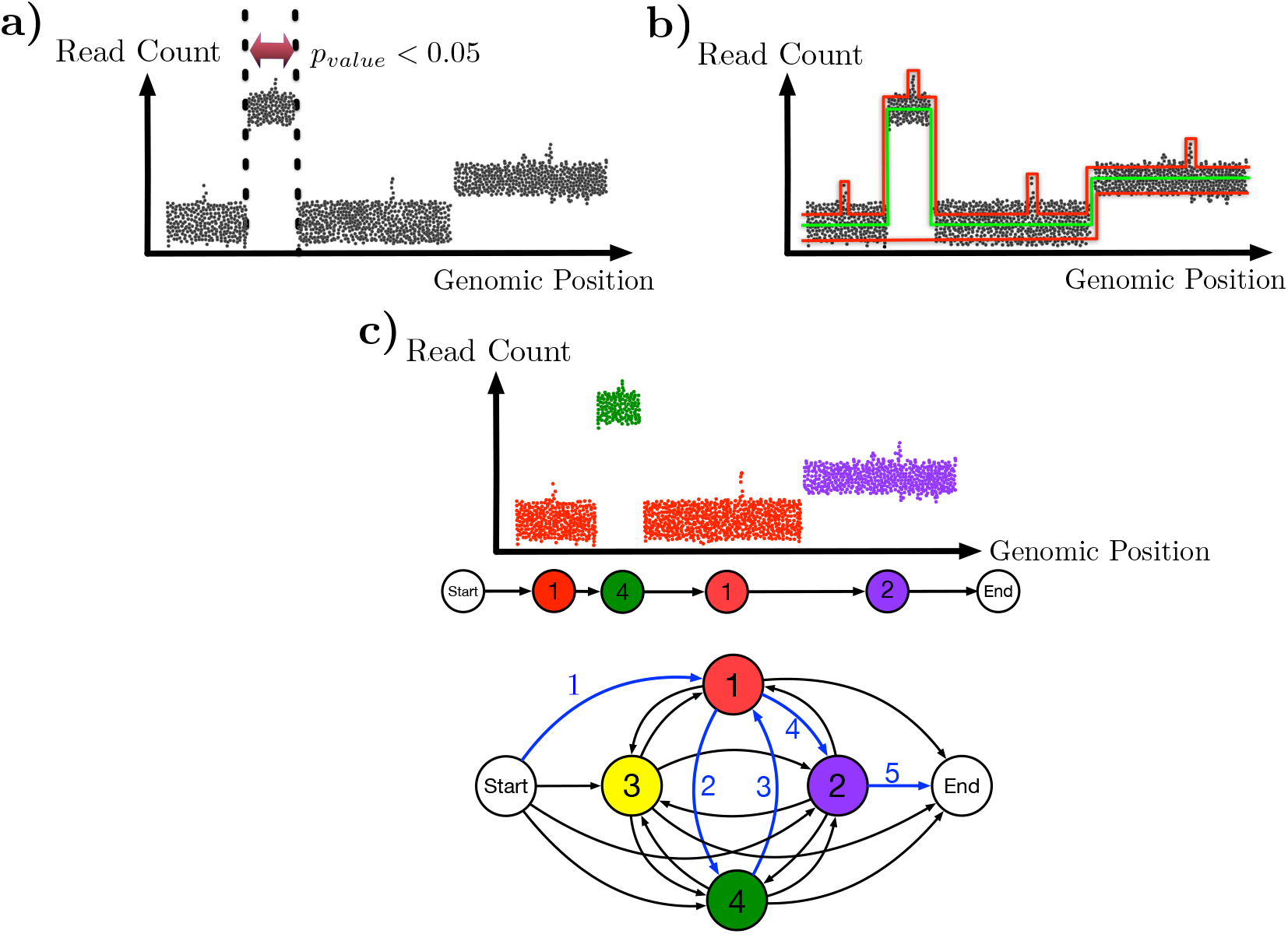
Illustration of the three approaches for segmentation. In all three panels, a scatter plot of the read count per bin with respect to the genomic position of the bin is shown. (a) The sliding-window approach. A window is passed across the genome and a genomic region within a window that is significantly different in terms of read count from the rest of the genome (e.g., the window defined by the two dotted vertical lines) is declared as a segment. (b) The objective-function approach. Three piecewise constant functions are shown (two in red and one in green) and represent segmentation candidates. Each piece in the function corresponds to a segment, and the value of the piece corresponds to the copy number at that segment. The function in green is the optimal one with respect to the fidelity to the data and constraint on the number of breakpoints, whereas the two in red are either over-segmented (top) or under-segmented (bottom). (c) The HMM approach. States of the HMM correspond to the different copy numbers and a transition between two different states indicates a change in the segment. In the read count panel, colors of the dots represent the absolute copy number of the various genomic bins (red for 1, yellow for 2, and green for 4) as obtained by parsing the data with respect to the HMM (bottom). The actual path of state transitions is shown in the middle and highlighted with blue arrows on the HMM as well. The arrows are numbered to indicate the order of the transitions.

### Calling the absolute copy numbers

Except for HMM-based methods, which identify the absolute copy number along with the segmentation, all other methods require a post-processing step to identify the absolute copy number for each segment after segmentation has been obtained. When DNA ploidy information is available, e.g., through flow cytometry data, the absolute copy number for a segment can be calculated by multiplying the genome-wide ploidy by the read count in that segment, divided by the mean read count across the genome [23]. When DNA ploidy information is not available, a copy-number multiplier is to be found which best approximates the normalized read counts in each bin to its closest integer number.

## CNA Detection Methods

Twenty-six methods for calling CNVs or CNAs are reviewed in this section, regardless of which platform the method was originally designed for. We particularly pay attention to and discuss how each method can be applied/generalized to single-cell DNA sequencing data. While copy number profiling is important beyond cancer, single-cell DNA sequencing holds great promise for deciphering intratumor heterogeneity [51], a confounding factor for diagnosis, prognosis, and treatment of cancer. Therefore, for each method, we focus on its potential applicability to tumor single-cell DNA sequencing data.

### Single Sample/Cell based Methods

#### HMMcopy

HMMcopy [61] is an HMM-based method that was originally designed for array CGH data. Several studies [34, 68, 83] applied it to large scale single-cell sequencing data. In particular, Vitak *et al.* [68] applied it to 16,698 single cells. HMMcopy uses a Bayesian network to model all parameters and hyper-parameters at once. Specifically, it explicitly models the case of outlier bins without penalizing them based on the neighboring bins. The original version of HMM-copy consisted of four states in the HMM (copy number loss, neutral, gain, and multiple gains), and was later extended to eleven states (zero to ten copies). It models the emission probability of each state by Gaussian distribution, whose mean and the inverse of variance are modeled by Gaussian and Gamma prior distributions, respectively. It models the transition probabilities by a Dirichlet distribution. The probability of a bin being an outlier is modeled by Beta distribution. HMMcopy estimates the state sequence and parameters simultaneously by blocked Gibbs sampling. Advantages of HMMcopy include its robustness to outliers and ameliorating the local minima problem during training, thanks to its explicit modeling of outliers as a state in the HMM and its usage of MCMC during training, respectively. Limitations of HMMcopy include its requiring manual calibration of more than ten parameters, as well as its inability to correctly estimate the ploidy, which leads to incorrect copy number estimation (as we demonstrate below).

#### SeqSeg

SeqSeg [15] detects CNAs in tumor-normal sample data. For each read from a tumor genome, the method maps it to a normal genome to identify its location *x* on the normal genome. Using the read data from the normal sample, left and right flanking regions of *x* each of which has a fixed number of reads *w* are identified; call them *l* and *r*, respectively. The method then counts the number of reads from the tumor genome that mapped to *l* and *r*, computes the ratio of these two numbers, and takes a statistically significant deviation from 1 as an indicating that *x* is a breakpoint between two regions with different copy numbers. Once the method is done with all reads, it merges adjacent breakpoints into single ones as long as the merged segment’s *p*-value is smaller than a threshold. Strengths of SeqSeg include a finer resolution than methods that rely on binning, robustness to GC bias by virtue of using *w*, and selectivity of somatic CNAs only. Limitations of SeqSeg include inapplicability of the method in the absence of a designated normal cell, and its inability to deal with CNAs at repetitive regions.

#### CNV-seq

CNV-seq [78] detects CNVs from tumor-normal sample data. It segments the genome into bins and transforms each bin’s read count ratio between the two samples from Gaussian ratio distribution to a standard Gaussian distribution by Geary-Hinkley transformation. For each bin, it checks its *p*-value for violating the null hypothesis of being copy number neutral. The optimal bin size is estimated by considering a desired *p*-value and the threshold of the read count ratio for calling a CNV, as well as the total number of reads for the two genomes. Strengths of CNV-seq include the adaptive bin size and that the sequencing bias is overcome by comparing two genomes. Limitations of CNV-seq include its inapplicability to tumor samples since it assumes the genome is diploid, its lack of scalability as it deals with one sample at a time, as well as its inability to identify the actual absolute copy number.

#### rSW-seq

rSW-seq [33] analyzes the read positions for tumor-normal paired samples to detect CNAs without using binning. Tumor and normal reads are tagged as positive and negative weights, respectively, and mixed together. The Smith-Waterman algorithm is applied to identify the region that has the maximal/minimal partial cumulative sum, which corresponds to copy number gain/loss, respectively. It then recursively applies the Smith-Waterman algorithm to the detected CNA regions for identifying overlapping copy number alterations. Advantages of rSW-seq include a finer resolution than methods that use binning. Limitations of the method include its inability to infer the absolute copy number, sensitivity to fluctuation in read coverage, and inapplicability in the absence of a designated normal cell.

#### FREEC

FREEC [12] automates the process of selecting the optimal bin size given a desired coefficient of variation. FREEC’s GC correction takes into account the copy number of each bin, different from most of the other methods which assume the copy number is the same during GC correction. Starting from estimating the polynomial regression from the bins with non-extreme GCs, it iteratively incorporates more bins, estimates their copy number and updates their polynomial regression for GC. FREEC also removes outlier bins by a mappability track, determined by the percentage of the non-N nucleotides in the reference genome, or the uniquely mappable positions in the bin. It uses a LASSO-based algorithm to segment the genome and computes the absolute copy number as the normalized read count multiplied by the given ploidy, rounded to an integer. Strengths of FREEC include its accounting for copy numbers during GC correction and its automated selection of the optimal bin size. Limitations of FREEC include its requiring the ploidy to be known, and that the convergence of GC correction’s algorithm highly relies on the accuracy of the initial guess of the polynomial regression.

#### CNAseg

CNAseg [29] is an HMM-based method that compares tumor-normal pair read count difference on NGS data. Before segmentation, it smoothes the low mappability region by a discrete wavelet transform, followed by removing the noisy wavelet coefficients in the frequency domain and inverse transforming the filtered signal to the space domain. CNAseg models the difference between tumor and normal read counts by a Skellam distribution. To avoid over-segmentation, CNAseg separates both normal and tumor samples into two groups. It then computes Pearson Chi-squared statistics for intra-type comparison (i.e., normal to normal, tumor to tumor), and uses its maximal value as the threshold for merging segmented bins in inter-type comparison. Advantages of CNAseg include its automated steps for removing low mappability region and choosing a threshold for merging segments. Limitations of CNAseg includes its inapplicability when no normal cell is provided.

#### ReadDepth

ReadDepth [49] detects CNAs without requiring the normal sample. It automatically estimates the optimal bin size given the read depth and a threshold of FDR. For segmentation, it models read count of each bin by a negative binomial distribution, the mean of which is the average read count across the whole genome, and the variance is tuned by a given variant/mean ratio value. Advantages of ReadDepth include its automated process of estimating optimal bin size and its applicability in the absence of a normal sample. Limitations include its assumption of the genome being diploid, its dealing with one sample at a time, and failing to identify the actual copy number.

#### CNVnator

CNVnator [1] uses a mean-shift algorithm to detect CNVs in a diploid genome. It pre-segments the genome by non-overlapping fixed bins, followed by GC correction. For each bin, it calculates the sum of the weighted difference between the neighboring bins’ read count and the read count of this bin. Two kernel density functions are used to model the read count difference and the distance between two bins. The bandwidth for the kernel density function for the read count difference is a function of the average read count, the read count of the bin of interest, and the overall standard deviation of the read counts. The bandwidth of the kernel density function for the distance between two bins, on the other hand, is gradually increased until a maximum value is reached, for the sake of detecting both large and small variations. In segmentation, CNVnator eliminates from consideration the bins whose read depths are significantly different from the neighboring bins. This step makes it possible to align two distant regions to the same copy number and thus be applicable to overlapping CNVs. It then uses a greedy algorithm to merge the segments, followed by calling a CNV through a statistical test. Advantages of CNVnator include its using mean-shift algorithm to incorporate both distance and read count difference between two bins, and its ability to align two distant regions to the same copy number by “freezing” the region in between. Limitations of CNVnator include heavily relying upon the diploidy of the genome for merging segments and calling CNVs, and its inability to identify copy number.

#### seqCBS

seqCBS [63] segments the genome by a generalized likelihood ratio test for every candidate breakpoint in a pair of tumor and normal samples. It uses a modified Bayesian Information Criterion (BIC) [85] to control the total number of breakpoints. The confidence interval is derived by a Bayesian framework considering both the read counts in an interval and the distance to the underlying breakpoints. seqCBS uses an Iterative Grid Scan to alleviate the computational burden. Strengths of seqCBS include its finer resolution in its bin-free strategy, as well as its ability to infer the confidence interval, which cannot be found in most of the other methods. Limitations of seqCBS include its inapplicability in the absence of a designated normal cell and that the absolute copy number is not identified.

#### seqCNA

seqCNA [50] corrects GC bias and filters out outlier bins, the steps to prepare for segmentation and copy number identification. Five filters are used to filter out outlier bins, including mappability, mapping quality, percentage of discordant read pairs, germline CNVs, and extremely high read count. In particular, a Wald-Wolfowitz test is used to select a threshold of a read count quantile to remove extremely high read count regions when the number of bins they span is relatively small. GC correction can be done in the absence or presence of a normal genome. When a normal genome is absent, seqCNA corrects read count first by a LOESS or polynomial regression. It then pre-segments the whole genome, and runs a second GC correction on only those large segments with the same copy number, large range of GC, and a small range of read count. Each segment undergoes a separate LOESS or polynomial regression. When a normal genome is present, seqCNA corrects read count based on the normal genome instead of the GC content. An advantage of seqCNA is its considering copy numbers when correcting GC bias. Limitations of seqCNA include the potential to over-filter outlier bins, usage of thresholds that are manually set, and its inability to correct GC bias on those smaller segments with smaller ranges of GC and larger ranges of read counts.

#### m-HMM

m-HMM [69] is an HMM-based method whose input is a target-reference genome pair, and it has been mainly applied to plant genomes. Its transition probability is a function of the distance between two bins. Its emission probability models the read count by a mixture Poisson distribution, the mean of which follows Gamma distribution. An Expectation-Maximization (EM) algorithm is used to estimate the parameters of the HMM. Advantages of m-HMM include its incorporating mixture Poisson distribution to tolerate read count fluctuations due to mapping and sequencing noise, as well as its modeling the distance between two bins in its transition probability. Limitations of m-HMM include its lack of actual copy number identification, and its dependence on the convergence of the EM algorithm for estimating the parameters (which could be an issue in practice).

#### Ginkgo

Ginkgo [24] is a web platform for analyzing single-cell copy number variations. It uses variable bin strategy to segment the genome into bins and a LOWESS regression for GC correction. To remove outlier bins, Ginkgo was applied to 54 normal diploid cells and a list of “bad bins” was created by testing the *p*-value of the normalized read count for each bin. It then uses CBS-based approach for segmentation, and identifies the absolute copy number by selecting the optimal multiplier to best align the normalized read count with an integer number. In addition to segmentation and calling absolute copy number, Ginkgo also clusters the cells, infers the phylogenetic tree among the cells, and provides plots to visualize GC bias and Lorenz curve, etc. Advantages of Ginkgo include its web-based platform and the post-processing steps such as phylogenetic tree inference and clustering of the cells. Limitations of Ginkgo include the fact that the list of bad bins is learned from a fixed set of normal cells, and its GC correction does not incorporate each bin’s copy number.

#### nbCNV

nbCNV [84] models the read count in each bin by a negative binomial distribution. The optimal bin size is selected when both Multiple Absolute Pairwise Distance (MAPD) and the approximation to the negative binomial distribution are considered. Segmentation is done in a Bayesian framework, in which the likelihood function is the approximation of the data to the negative binomial distribution, and the prior probability models smoothness of the CNA profile. Numerical solution for maximum a posterior probability estimate is computed through the framework of Alternating Direction Method of Multipliers (ADMM), while fixing the over-dispersed parameter by an MLE-based method. nbCNV is advantageous in applying Bayesian framework and its numerical solution to single-cell and its automated process of selecting the optimal bin size. It is limited in not being able to pool cells together for segmentation, and thus not taking advantage of the fact that the cells may share breakpoints.

#### AneuFinder

AneuFinder [5] is an HMM-based method. It uses the variable-size bin strategy so that the read count of each bin are the same and the value of the read count in each bin is obtained from fixed-sized bin strategy. It then corrects GC content and does quality control using a multivariate clustering approach implemented in mclust [20–22] which combines several metrics defined in [5]. For each bin, there are eleven states, representing copy numbers from 0 to 10. Except for copy number 0 that is modeled by delta distribution, all others are modeled by negative binomial distribution. Parameters of the HMM are learned using the Baum-Welch algorithm [7]. The copy number state that has the highest posterior probability is then selected to be the copy number for this bin (i.e., it uses posterior decoding). Advantages of AneuFinder include its quality control for single-cell sequencing. Its limitations include its failure to correct GC bias according to the copy number that each bin has.

#### SCNV

SCNV [73] is an extension of seqCBS to single-cell sequencing. It automatically identifies normal cells and pools them to serve as the normal sample pair. Such identification is made by segmenting the genome by CBS with fixed-size bin, followed by computing the L2 norm of the difference between the median read count of each segment and the average read count of the whole genome. This step also serves to remove outlier cells. SCNV then segments the genome by pairing up each tumor cell with pooled normal cells and adopting the strategy of seqCBS for segmentation. The threshold of segmentation is calibrated by running SCNV on (pooled) normal cells. This is based on the assumption that all cells are sequenced by the same platform and have similar error profiles. To identify the absolute copy number of each segment, SCNV infers the ploidy by modeling it as a damped sine wave function. The rationale is similar to finding the ploidy that minimizes the squared sum of the difference between the estimated copy number and its nearest integer, as used in Ginkgo [24]. Strengths of SCNV include its automated process of identifying normal cells, outlier cells, tumor cell ploidy, and the threshold for segmentation. Limitations of SCNV include its inability to segment cells by pooling them together, which makes it fail to take advantage of the fact that cells may share breakpoints.

#### CNV IFTV

CNV IFTV [81] uses an isolation tree strategy to calculate an anomaly score for each bin, followed by merging the neighboring bins by the anomaly score to overcome normal contamination (the presence of normal cells in tumor sample). The anomaly score incorporates the difference of the read count in each bin in rank by comparing the path length from the root to this bin in the isolation tree with the average path length of other bins. It then smoothes the anomaly score by total variation, calculates the *p*-value of the anomaly score on tentatively merged bins, and identifies those bins having small *p*-values. Advantages of CNV IFTV include its robustness to GC content, mappability, and the bias of duplication to deletion, thanks to its use of the read count rank rather than the value itself. Limitations of CNV IFTV include its inapplicability to tumor samples since the identification of the absolute copy number assumes the ploidy being 2.

### Multiple Sample/Cell based Methods

#### PennCNV

PennCNV [70] is an HMM-based method that detects germline CNVs at a kilobase-resolution using SNP genotyping data. The transition probability is state-specific and distance-dependent. The parameters are learned by the Baum-Welch algorithm [7]. Advantages of PennCNV include that it incorporates family information, the Mendelian inheritance rule based on CNVs inferred from parental SNP genotyping data, to more accurately infer CNVs. Limitations of PennCNV include its inapplicability to single-cell sequencing, as it was originally designed for SNP genotyping data, as well as its modeling of up to two-copy gain, making it not suitable for higher-copy gain detection.

#### RDXplorer

RDXplorer [80] divides the genome into fixed-sized bins, and approximates the GC-corrected read count in each bin as a sample from a normal distribution. For each bin, it then tests to see if its upper-tail or lower-tail z-score is larger or smaller than a pre-defined score that takes into account the size of the region of interest and an FDR threshold. It then iteratively merges the neighboring bins and does the same test on the merged bins until no more mergers can be made. To detect polymorphic CNVs for multiple samples, it makes a pairwise comparison between the read count of each sample and that of the rest of the samples by t-test. If at least one of the tests has a *p*-value smaller than a threshold, and its median read count differs from the rest by more than a threshold, this CNV is designated as polymorphic. Since RDXplorer assumes the whole genome, except those duplicated or deleted regions, is diploid, it is not applicable to aneuploid tumor samples. Furthermore, it only infers whether a region has a deletion or duplication, but does not infer the actual copy number. RDXplorer’s strategy to identify polymorphic CNV, however, is applicable to single-cell sequencing data. But extra work is needed to normalize raw read count to read count ratio, since read coverage for each single cell is not necessarily the same. However, the large number of cell-to-cell comparisons render this strategy for detecting CNAs limited to very small data sets.

#### cn.MOPS

cn.MOPS [80] uses a Bayesian framework that pools all samples’ read counts together for segmentation. Assuming a Poisson mixture distribution, it uses an EM algorithm to estimate the posterior of two parameters: *α*, the percentage of each copy number, and *λ*, the average number of read counts per bin for copy number 2. It then estimates the information gain compared with the prior of *α* which peaks at copy number 2. Segmentation is done by merging adjacent bins based on the estimation of information gain from all samples, followed by a posterior inference of the best fitting absolute copy number. Advantages of cn.MOPS include its usage of information gain and mixture Poisson distribution, and its design for segmentation on multiple samples and thus is generalizable to single-cell sequencing. One limitation of cn.MOPS is its assumption that the copy numbers peak at 2.

#### The method of Zhang et al

Zhang et al. [86] extended the CBS algorithm [55] to multiple samples, with an addition of the equations for inferring the significance levels and the algorithms to infer the samples that carry the inferred CNA. Advantages of this method include its pooling all samples/cells together for segmentation, and its ability to infer the samples carrying the CNA. Limitations include its inability to infer the absolute copy number and that it was originally designed for probe data instead of sequencing data.

#### BICseq

BICseq [77] models each tumor/normal read by a Bernoulli trial, and that the probability of sampling a tumor read out of the mixed samples is a piecewise constant variable. The change of such piecewise constant variable is where the breakpoints are. Using BIC, it manages to control the number of breakpoints. In a bottom-up manner, it greedily merges the neighboring bins that may result in the lowest BIC value. It then assigns the copy ratio and significance to each segmented region, as well as the confidence interval. One advantage of BICseq is its modeling the read count by Bernoulli trial, avoiding using Poisson or negative binomial distributions which do not necessarily approximate the distribution well [77]. Limitations of BICseq include its inapplicability when a normal paired sample is not available, and its assumption that the genome is generally copy number neutral.

#### JointSLM

JointSLM [44] uses an HMM to jointly segment multiple genomes simultaneously. Advantages of JointSLM is its pooling all samples together for segmentation. Limitations of JointSLM include that all samples have to adopt the same copy number changes, its assumption of the diploidy of the genome, and its fixed prior probability of a breakpoint.

#### GFL

GFL [87] simultaneously segments multiple genomes on SNP array data by minimizing an objective function, which includes a total variation and a generalized fused lasso for controlling the change points in a group of samples. It then uses an MM [37] framework that adopts a surrogate function in each iteration for finding the solution to this optimization problem. Advantages of GFL include that it is computationally light thanks to using the MM framework, and its ability to detect CNVs occurring to a small number of samples from a set of sample data. One limitation of GFL is its inapplicability to single-cell sequencing as it was originally designed for SNP array data.

#### CopyNumber

CopyNumber [53] uses Piecewise Constant Fitting to segment single-sample, multiple-sample, or allele-specific array-CGH data while being constrained by the total number of segments. It first reduces the effect of the outliers by winsorizing the extremely high/low signals, and then uses a dynamic programming approach to identify the breakpoints quickly. Advantages of CopyNumber are its pooling all samples/cells together for segmentation, and it is computationally light. Limitations of the method include its failure to model single-cell specific error profiles and its inability to assign the absolute copy numbers.

#### MSeq-CNV

MSeq-CNV [45] combines both read depth and discordant read pairs and jointly estimates the absolute copy number for multiple samples. It models the read depth at each bin for each sample as a Poisson distribution, and the read pairs that have the insert size consistent with the clone library insert size a binomial distribution. The two pieces of information are independently modeled, and the weight of the number of samples having each copy number has a Dirichlet prior distribution. An EM algorithm is used to estimate the parameters, initialized in a way to give more weight on copy number 2, assuming most of the bins in most of the samples are diploid. Advantages of MSeq-CNV include its inclusion of discordant read pairs in inferring CNAs and its applicability to multiple samples. Limitations of the method include its inapplicability to single-cell sequencing since the signal of discordant read pairs are not expected to be often observed in the low coverage data or in chromosomal amplifications.

#### SCOPE

SCOPE [72] jointly segments all cells by a generalized likelihood ratio test proposed originally in [86] and later adopted by seqCBS [63] and SCNV [73]. It then uses a modified BIC [85] to control the number of segments, the same strategy adopted by [63] and [73]. In the normalization step, SCOPE adopts a Poisson latent factor model that considers the cell-specific GC content bias, cell-specific library size, bin-specific read depth, the absolute copy number of the bin of interest, and the Poisson latent factors modeling other systemic biases. Such normalization is a generalization from CODEX2 [31], a CNV detection tool for whole-exome and targeted sequencing data. Since normal cells are required as negative controls in normalization, it identifies normal cells by the Gini coefficient of each cell. The outlier cells are also identified by the Gini coefficient in this step. Advantages of SCOPE include its pooling all cells together for segmentation, considering the copy number for GC correction, utilizing bin-specific read depth variable for normalization, and its considering latent factors that are bin- and cell-specific. Limitations of SCOPE include that its normalization step requires at least 10 to 20 normal cells present in the pool, and that its joint segmentation does not consider lineage among the cells.

## Performance of Three Representative Methods

To better understand the strengths and limitations of current approaches for CNA detection from single-cell DNA sequencing data, we selected three methods that have been commonly applied to cancer single-cell genomic data and represent the three segmentation approaches: HMMcopy [61] uses the HMM-based approach, Ginkgo [24] uses the sliding window approach, and CopyNumber [53] uses the objective function based approach.

We designed three experiments to evaluate the performance of the three methods under different conditions. The first experiment was designed to evaluate the recall and precision of the CNA detection methods. We designed the simulation study in this experiment to produce single cells that have ideal read count variability and normal ploidy levels ranging between 2 and 3, so that we can learn how the methods perform when the data is relatively not challenging. The second experiment was designed to evaluate the performance of each method under a variety of ploidy levels. Specifically, we simulated single cells whose ploidies range from 1.5 to 5.26 (the ploidy of a cell is defined as the average copy number across the cell’s genome). We then compared the recall and precision of the three methods on the simulated data at different ploidies. The third experiment was designed to assess the performance of each method under different coverage variabilities. In particular, we simulated single cells whose coverage variabilities mimic those produced by three single-cell sequencing technologies (MALBAC, DOP-PCR and TnBC) [76] that have been used for CNA detection.

### Precision and recall of the methods

In the first experiment, we simulated the evolution of 10,000 cells from which we generated, through sampling without replacement, three 1000-cell datasets. For each cell, we simulated read data using a simulator that we developed (see the “Methods” section). We then aligned the reads back to the reference genome using BWA with default parameters [38, 39]. Finally, we ran each of three methods on the resulting BAM files, and computed the recall and precision of each method based on the ground truth generated by the simulator.

We assessed the methods’ performances in coarse- and fine-grained analyses. For the coarsegrained evaluation, we inspected the breakpoint positions as well as whether they were consistent with the ground truth in terms of estimated gain/loss state (rather than the actual value) in the copy number. The predicted breakpoint is counted as consistent with the ground truth whenever it has the same status (i.e., the copy number increases or decreases) and its genomic location is within a certain distance of its counterpart in the ground truth. We varied the value of this distance to study the methods’ accuracies. Each ground truth breakpoint was matched by at most one predicted breakpoint to avoid double counting of the true positive calls. For each method, we varied a parameter to obtain the receiver operator characteristic (ROC) curve, the details of which are described in the caption of fig. 3.

**Figure 3:**
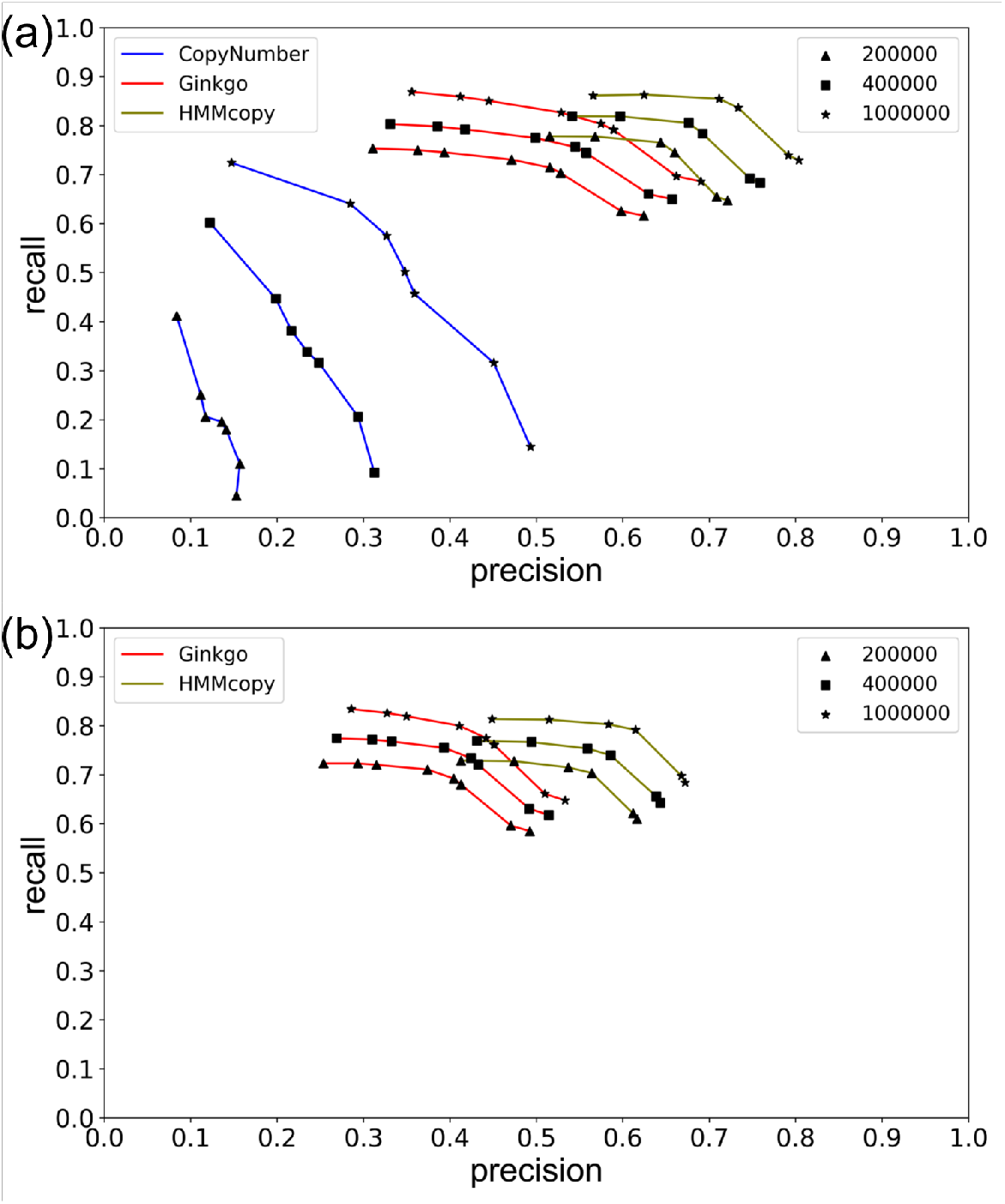
ROC curves of the three methods HMMcopy, Ginkgo, and CopyNumber. (a) Coarse-grained analysis results, and (b) fine-grained results. For each method, the results based on three thresholds of correctness are plotted. For HMMcopy, *nu*, which controls the suggested degree of freedom between states, was tuned to take on the values 0.01 (rightmost), 0.1, 2.1 (the tool’s default), 4, 10, and 20 (leftmost). For Ginkgo, *alpha*, which controls the significance level to accept a change point, was tuned to take on the values 1e-1000 (rightmost), 1e-100, 1e-10, 1e-5, 1e-4, 1e-3, 1e-2 (the tool’s default), 0.02, and 0.05 (leftmost). The dots corresponding to values 1e-5 and 1e-10 in coarse-grained analysis overlap. For CopyNumber, *gamma*, which is the weight of the penalty on changing a state, was tuned to take on the values 40 (rightmost, and the tool’s default), 10, 5, 4, 3, 2, and 1 (leftmost).

A preliminary analysis of CopyNumber on the data revealed that the method achieves extremely low recall and precision. Since CopyNumber pools all cells together for breakpoint detection, we suspected that this poor performance owes mainly to the method’s lack of sensitivity in detecting breakpoints shared by only a small number of cells. Therefore, to allow for more meaningful comparison of CopyNumber to the other two methods, we eliminated breakpoint pairs shared by fewer than five cells in the ground truth and used the resulting new ground truth to evaluate CopyNumber’s recall and precision (but we did not filter the breakpoints for the other two methods). As fig. 3a shows, CopyNumber still has, by far, the poorest performance. We hypothesize that for a breakpoint to be detectable by CopyNumber, it needs to be shared by a large number of cells. We further checked this by calculating the number of cells sharing a breakpoint that is called or missed by CopyNumber, and found that there is a significant difference between the two sets (*p*-value *<* 9.019e-09 for Student’s t-test with mean 9.27 versus 5.35). We also observe that as the tolerance threshold for the detected breakpoint position increases, improvement in Copy-Number’s performance is much larger than the improvement in the performance of the other two methods. However, even with the most forgiving threshold (two breakpoints are considered to be the same if their positions are within 1 million basepairs of each other), CopyNumber still has poorer performance than the other two methods even under the most stringent threshold. Overall, the results in fig. 3a show that (1) HMMcopy generally outperforms the other two methods, with Ginkgo being a close second, and (2) that even HMMcopy’s best recall and precision are around and 0.7, respectively.

In the fine-grained analysis, we focused on the agreement of the absolute copy numbers on both 5’ and 3’ of an inferred breakpoint with those of the ground truth, in addition to the requirements on gain/loss consistency and distance. Since CopyNumber does not predict the absolute copy numbers for an individual cell, it is not considered in this analysis. Surprisingly, HMM-copy’s prediction of the absolute copy number is not stable, leading to a bimodal distribution of both recall and precision (Additional file 1: Figure S1). We found that cells with low recall and precision correspond mainly to cases where HMMcopy made inaccurate estimates of the cells’ ploidies (Additional file 1: Figure S2). We then selected only those cells for which the ploidy was correctly predicted (i.e., 2 or 3), and plotted the ROC curve of HMMcopy on them. We found that HMMcopy performed generally better than Ginkgo (fig. 3b), which is in agreement with the coarse-grained analysis. The recall and precision for the two methods dropped, which is expected since the true positives and negatives are now measured most stringently. However, we observed that the difference in results between the coarse- and fine-grained analyses is not large, suggesting that once the breakpoint is found by these methods, predicting the absolute copy number can be attained quite accurately. This is especially true for Ginkgo whose ploidy prediction is stable.

Similar results were observed on the other two datasets (Additional file 1: Figures S3 and S4). The results in fig. 3 were obtained under default parameters except for the parameters that were tuned to generate the ROC curves (*alpha*, *gamma*, and *nu*). However, we found that the value of parameter *strength* in HMMcopy has to be much larger than the default value in order to make the results more expected, i.e., increasing recall is accompanied with decreasing precision, and vice versa. We therefore set *strength* to be 10 million. According to HMMcopy’s users’ guide, *strength* is the parameter that controls how much the initial values of *e*, which controls the probability of extending a segment, remains the same throughout the iterations. We found that setting *strength* would help to have a good quality control of the result by making the initial setting of *e* last throughout all the iterations. Apart from parameters *nu* and *strength*, we found that *e* is also an important parameter in HMMcopy. The larger the value of *e*, the smaller the chance that the breakpoint is detected. To explore which combination can yield the best performance for HMMcopy, we varied both *e* and *nu* and calculated the F1 score. The performance of HMMcopy is the best when *nu* is 4 and *e ≥* 0.999999 (Additional file 1: Figure S5).

We also analyzed the computational requirements in terms of running time and memory consumption for the three methods on a 1000-cell dataset (fig. 4). Ginkgo is the slowest among the three and CopyNumber is the fastest. HMMcopy is in between Ginkgo and CopyNumber in terms of running time. For memory consumption, Ginkgo is the most demanding of the three, followed by CopyNumber. HMMcopy is the lightest in terms of memory consumption. Note that in running Ginkgo, we eliminated the steps of generating figures such as heatmaps and copy number profile, so that these do not affect the running time and memory in comparison with the other two methods. For CopyNumber, an extra step is required to generate its input file. We used the intermediate result of HMMcopy, i.e., the GC corrected read count on each bin, as the input to CopyNumber. We take the time for calculating this intermediate file into account for CopyNumber for a fair comparison. Since CopyNumber processes all cells together, we suspect that more cells will require more memory, whereas Ginkgo and HMMcopy’s memory requirements are not affected by the number of cells involved.

**Figure 4:**
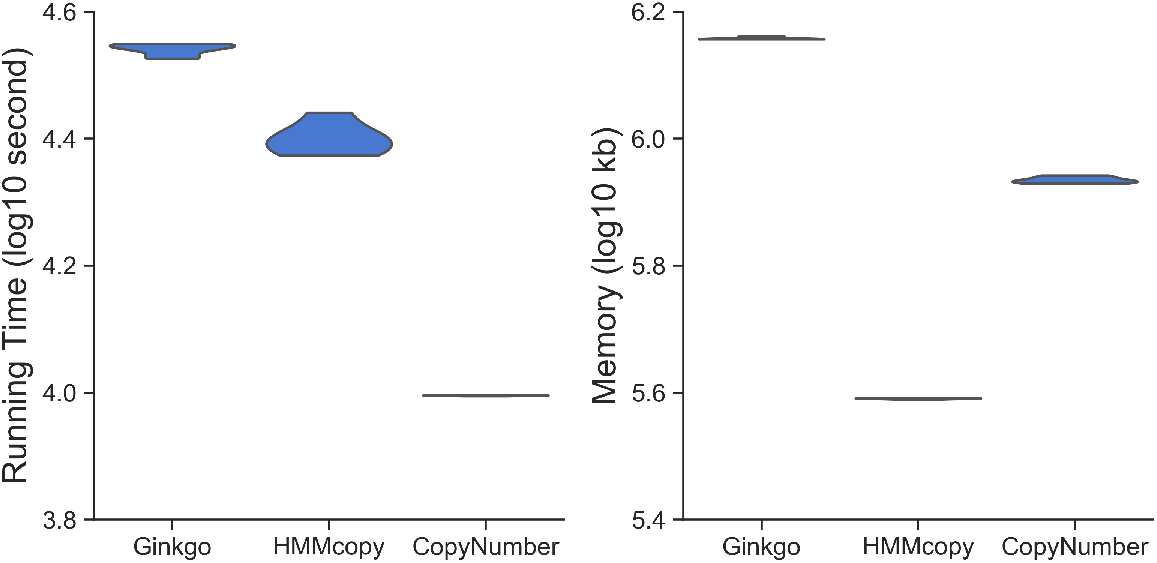
Computational requirements of Ginkgo, HMMcopy, and CopyNumber. Results are for analyzing a 1000-cell dataset on Intel(R) Xeon(R) CPU E5-2650 v2 whose clock speed is 2.60GHz. Left and right panels correspond to running time (in log10 of seconds) and memory consumption (in log10 of kb). The running time and memory were recorded for using different parameters as described in fig. 3. As Ginkgo’s running time increases more than twofold for *α* = 0.05, we treated it as an outlier and did not include this running time point in this plot.

In summary, we found that HMMcopy is the most accurate in predicting breakpoints among the three. When HMMcopy’s prediction of ploidy is accurate, its recall and precision of predicting the absolute copy number are the best among the three methods. However, it is not as stable as Ginkgo in terms of absolute copy number detection since its prediction of ploidy is wrong for many cells (49.4% for default values of *nu* and *e*). CopyNumber’s recall and precision are the worst of the three methods. Moreover, it cannot predict the absolute copy number for each individual cell, and thus is not as applicable as the other two methods.

### The effect of ploidy on performance

To test the robustness of the methods to different ploidies, we varied the ploidy by tuning the parameters that control whole chromosomal amplifications and the rate of deletion (see the “Methods” section). We varied the ploidy from 1.5 to 5. Specifically, we used 1.5, 2.1, 3, 3.8, and 5.26 (we refer to them as 1, 2, 3, 4, and 5, respectively, hereafter), and generated three datasets for each ploidy. We tuned the coverage parameter for each ploidy so that the total number of reads for different ploidies are approximately the same to avoid adding reads for larger genomes resulting from higher ploidies.

We ran each method using their default parameters (except the *strength* parameter in HMM-copy). Finding CopyNumber’s recall to be zero using the default *gamma*, we tuned *gamma* using the optimal value, i.e., 5, shown in fig. 3). We then found the recall greatly increased with this setting. Similar to the previous experiment, we again removed those breakpoint pairs shared by ≤ 5 cells from the ground truth for evaluating CopyNumber’s performance.

We used different combinations of the parameters to simulate high-ploidy cells (details are in the “Methods” section), i.e., 4 and 5, and found that in the absence of odd and intermediate copy numbers, HMMcopy’s inference of the ploidy and absolute copy numbers were inaccurate (Additional file 1: Figure S6). This is also the case for Ginkgo in the absence of the odd copy numbers. However, despite the lack of intermediate copy numbers, Ginkgo correctly predicted absolute copy numbers for the case of ploidy=5, showing that Ginkgo is more robust to changes in ploidy than HMMcopy in terms of predicting absolute copy numbers. In summary, the lack of odd or intermediate copy numbers in the data led to wrong predictions of absolute copy numbers. We then tuned the simulator’s parameters so that in higher ploidies there are odd and intermediate copy numbers to avoid the extremely hard cases for CNA detection (details are in the “Methods” and “Conclusions” sections). fig. 5 shows the precision and recall for the three methods for coarse- and fine-grained analyses, respectively.

**Figure 5:**
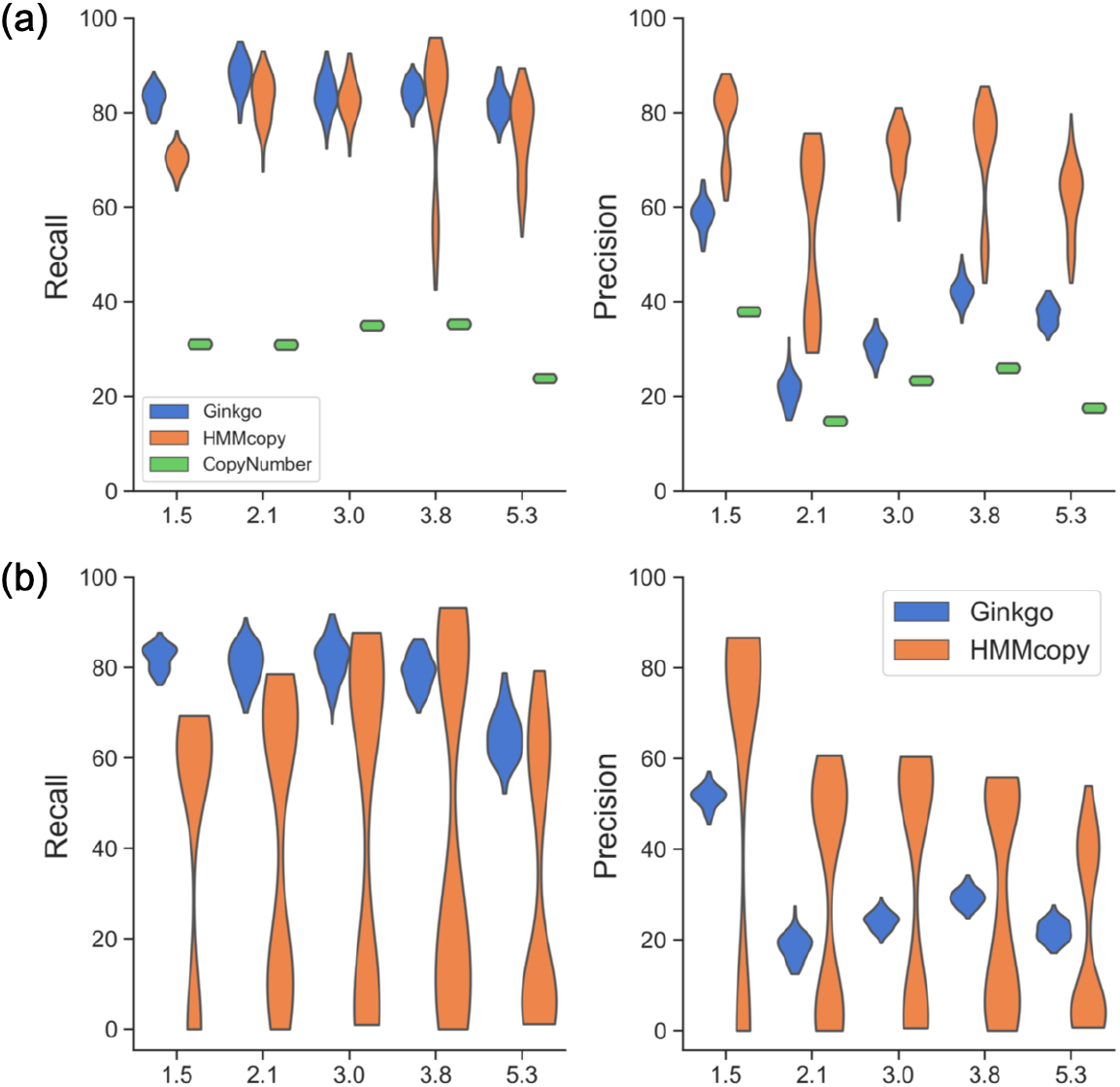
Recall and Precision of Ginkgo, HMMcopy, and CopyNumber as functions of the ploidy. The ploidy level is varied and the results are based on the (a) coarse-grained and (b) fine-grained analyses. The ploidies of the simulated data were 1.5, 2.1, 3.0, 3.8, and 5.3.

In the coarse-grained analysis, Ginkgo’s recall is *>* 0.8 in general, but its precision is relatively low (i.e., *<* 0.4) for ploidy 2 and 3. This is probably because it was affected by the variability of the read counts and over-segmented the genome. With similar recall, HMMcopy has higher precision at all ploidies. CopyNumber’s recall and precision are low (*<* 0.4) for all ploidies, with low recall and precision for ploidy 5 and low precision for ploidy 2.

In the fine-grained analysis, Ginkgo’s recall and precision dropped by about 10% as compared with the coarse-grained results. Its recall dropped the most for ploidy 5, indicating the challenge in accurately predicting the absolute copy number for high-ploidy cells. Although the odd and intermediate copy number states are present in this simulated data, HMMcopy’s precision and recall are still bimodally distributed for all ploidies. As we observed cells whose incorrect ploidy prediction led to wrong prediction of absolute copy numbers in the previous experiment, these bimodal distributions further showed that such wrong prediction can widely occur to cells with different ploidies.

Similar results were observed on the other two datasets (Additional file 1: Figures S7-S10).

### The effect of coverage on performance

To evaluate the performance of each method under different single-cell sequencing technologies, we sampled 20 cells from the simulated dataset and simulated their sequencing at four levels of coverage variabilities, corresponding to MALBAC, DOP-PCR, TnBC and Bulk sequencing (see details in the “Methods” section) and ran the three methods on each of them. We generated three datasets for each level of variability. fig. 6 show the performance on one of the datasets. With decreasing variability, all three methods’ recall increased under both the coarse- and fine-grained analyses. Ginkgo’s and HMMcopy’s precisions increased with decreasing variability. CopyNumber’s precision, on the other hand, stays the same regardless of the coverage variability level, whereas its recall generally increases. In summary, better sequencing technology leads to better performance. The best that can be ever obtained (Bulk sequencing) is about 15% higher than the worst (MALBAC) for recall, and slightly higher for precision.

**Figure 6:**
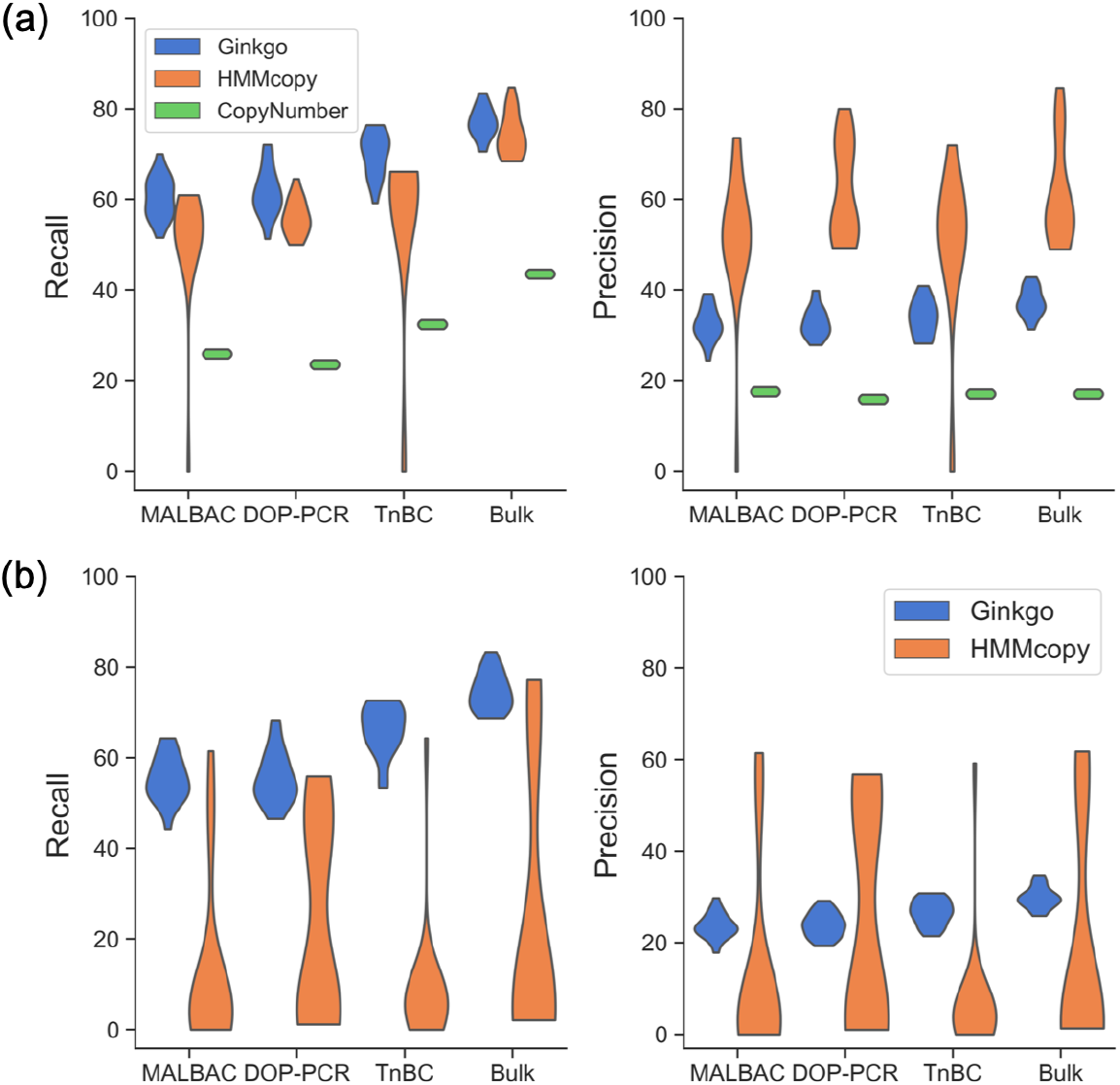
Recall and Precision of Ginkgo, HMMcopy, and CopyNumber as functions of the coverage. The coverage is varied and the results are based on the (a) coarse-grained and (b) fine-grained analyses. The coverages are varied to mimic those produced by MALBAC, DOP-PCR, TnBC and Bulk sequencing.

We looked into the copy number profiles in cases where HMMcopy’s precision and recall were effectively 0 (one such case is illustrated in Additional file 1: Figure S11). We found that choosing a wrong ploidy from the set of candidate ploidies by HMMcopy may result in a copy number profile in which all the segments are predicted to have the same absolute copy number, whereas the closest profile to the ground truth is among the reported non-optimal results. We observed that in such cases, the wrong choice of ploidy may affect both the segmentation and inference of the absolute copy number of those segments.

Similar results were observed on the other two repetitions (Additional file 1: Figures S12-S15).

### Performance on a real dataset

In real data analysis, due to the lack of ground truth, we evaluated the performance of the three methods in two ways. First, we evaluated the consistency among the three methods. The more overlap among the methods, the more consistent they are. Second, we inferred a maximum parsimony tree using PAUP [65] from the inferred copy number profiles and calculated the smallest number of copy number changes for each bin across all branches of the tree, where the genome at the root of the tree is assumed to be diploid. The rationale for the latter way of assessing performance is that if the CNAs called by a method result (under a parsimony analysis) in a very large number of changes of the copy number at any bin, then one plausible explanation is error in the calls (another plausible prediction is that, for some reason, the locus corresponding to that bin has repeatedly gained and lost copies during the evolution of the cells which could be indicative of some interesting biological processes at play).

We downloaded single-cell DNA sequencing data of seven samples (the median number of cells in the seven samples is 47) from [32] and selected those pre-treatment samples whose CNA profiles had not changed due to treatment compared with mid-treatment and post-treatment ones. We then ran the three methods with default parameters (except for the *strength* parameter in HMMcopy, as discussed above) on the single cells in each sample.

For assessing accuracy, we generated for each sample a Venn diagram of the predictions based on all three methods, where predictions by two methods were deemed consistent according to the same rule we used in the simulation study (in assessing consistency between predictions and the ground truth). fig. 7a shows the results for Sample 102 (Additional file 1: Figures S16-S18 show the results for the other six samples). It can be seen that 47% of Ginkgo’s calls overlapped with the other two methods, leaving a large portion as unique calls. HMMcopy overlapped well with the other two methods, with 22% of unique calls. In particular, HMMcopy overlapped well with Ginkgo: 76% of HMMcopy’s calls overlapped with Ginkgo. CopyNumber’s overlap with Ginkgo was larger than that of HMMcopy (65% versus 43%). The overlap among the three methods is a very small portion of the union of all calls (8%), indicating a very large inconsistency among the three methods. From these results, we observe that HMMcopy performed the best among the three in breakpoint calling, if we consider consistency with other methods as a metric of quality, which is consistent with what we observed on the simulated data.

**Figure 7:**
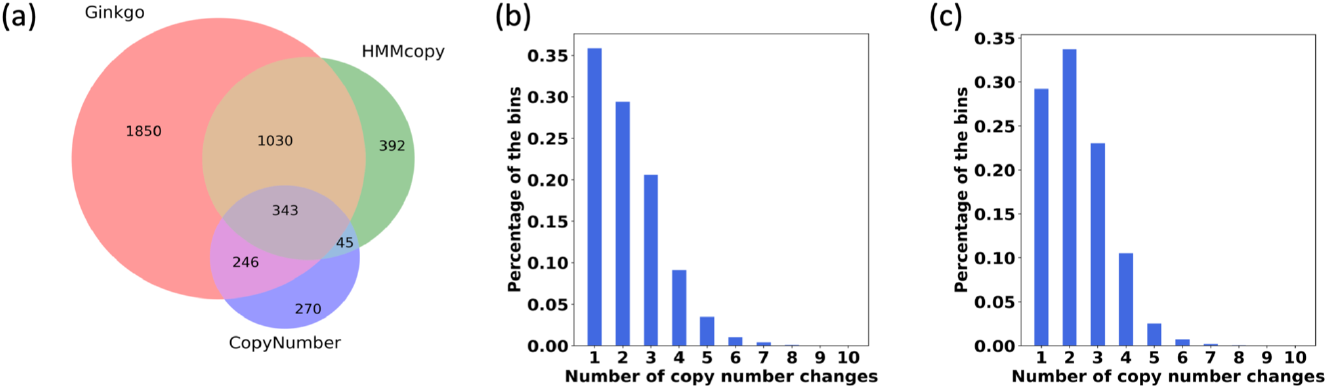
Comparison of HMMcopy, Ginkgo and CopyNumber on Sample 102 in [32]. (a) Venn diagram of the breakpoints from Ginkgo, HMMcopy and CopyNumber. Breakpoints from two methods are counted as overlapping if they are within 400,000bp of each other. (b) Distribution of the copy number changes (under a parsimony analysis) per bin based on the copy number profiles obtained by HMMcopy for the seven samples. (c) Distribution of the copy number changes (under a parsimony analysis) per bin based on the copy number profiles obtained by Ginkgo for the seven samples. For (b) and (c), a maximum parsimony tree was inferred from the copy number profiles of the cells, and the minimum number of copy number changes per bin along all branches of the tree was computed by parsimony analysis. The percentages of bins with each number of copy number changes are plotted.

We then investigated the smallest number of changes required to explain the copy numbers detected by Ginkgo and HMMcopy (again, CopyNumber does not detect absolute copy numbers, which is why it is excluded in this analysis). fig. 7b and fig. 7c show the distributions of copy number changes based on HMMcopy and Ginkgo’s results, respectively (Additional file 1: Figures S17-S18 show results for the other six samples). Interestingly, four out of seven samples (samples 102, 132, 152, and 302) showed a higher number of bins that have one copy number change than ones with no copy number changes. The other three samples (samples 126, 129, 615) have the most bins that had no copy number changes at all. Generally the number of bins that had copy number changes decreased with the increasing number of changes. On the other hand, based on the HMMcopy results, all samples showed much higher percentage of no copy number change than those with some copy number change.

## Conclusions

Single-cell DNA sequencing data holds great promise for elucidating the complex biological processes that underly human diseases, including cancer. Copy number aberrations have been implicated in cancer, and their accurate detection from single-cell DNA sequencing data is of great potential to diagnosis and treatment of cancer. In this paper, we reviewed 26 methods that have been developed for copy number variation/aberration detection, but not necessarily specifically for single-cell data. In our review, we outlined the seven general steps that a CNV/CNA detection pipeline follows, and discusses the 26 methods in light of these steps. For each method, we highlighted its advantages and limitations, paying special attention to applicability of the method to single-cell DNA sequencing data from cancer genomes. Two important outcomes of CNA detection are segmentation of the genome into non-overlapping regions with different copy numbers and inference of the actual copy numbers of the different segments. We reviewed the three different approaches to segmentation, and discussed for each of the 26 methods which of these two outcomes the method is capable of producing.

We then selected three methods that represent the three different segmentation approaches and that have been widely applied to single-cell data and benchmarked them on simulated as well as real data sets. The methods are CopyNumber, Ginkgo, and HMMcopy. We compared the three methods on simulated data generated under different settings that reflect varying degrees of complexity in the data. To accomplish this task, we developed a simulator that is flexible to simulate different scenarios and also mimic realistic data. We found that HMMcopy performs the best for breakpoint detection. However, HMMcopy is not stable in inferring the absolute copy number. Ginkgo performs well for both breakpoint detection and inference of the absolute copy number. CopyNumber is not as sensitive as the other two methods. We also looked into the performance of the three methods when ploidies were varied. We found that data with higher ploidies presented challenges for Ginkgo. HMMcopy is the most robust in terms of breakpoint detection among the three methods regardless of the ploidy, but its inference of the absolute copy number is not accurate for all ploidies. Both recall and precision of CopyNumber are the worst among the three methods. To explore the effect of technology artifacts on the accuracy of the methods, we simulated data that mimics the variability in coverage corresponding to MALBAC, DOP-PCR, TnBC, and Bulk. We found that all three methods’ recall generally increases with the improvement in the technology, with smaller observed change in their precision. We then applied the three methods to real data and evaluated their performance by analyzing the shared and unique detections they made as well as counting the total number of copy number changes must be invoked based on their detections. We found a good amount of overlap in detections between Ginkgo and HMMcopy. We also found that HMMcopy’s detections result in fewer copy number changes than Ginkgo’s.

Our review of the 26 methods and benchmarking of the three representative methods highlights several points. First, the ploidy and coverage of the genome under analysis affect the ability of a computational method to detect CNAs. Second, there is much more power in analyzing all cells in a data set simultaneously, as the fact that they all evolved from a common ancestral cell provide not only signal for the inference, but also accounts naturally for model complexity and regularizes the number of changes in copy numbers. Third, using algorithms and tools from the field of phylogenetics can help significantly in this area. As we demonstrated above on the biological data set, parsimony analysis of phylogenetic trees can be used to identify regions with large numbers of changes in the copy number, especially convergent changes, and those can be further inspected for determining whether the high rate of change reflects a biological process that is worth pursuing or due to sequencing or computational inference error artifacts. Finally, our results show that while existing methods are a good step in the right direction, there is need for developing more accurate methods for CNA detection, especially ones that are designed specifically to model the specifics of single-cell DNA sequencing.

## Methods

### Simulation

Two steps are involved in simulating reads for single-cell sequencing. First, the cell tree is generated, where the nodes are the cells, and the edges represent the parent-daughter cell relation. The leaf nodes represent the single cells that are sampled from the patient; the internal nodes represent the cells that existed in the past and were not sampled. We set the root node as a normal genome without any CNA, assuming that all CNAs are somatic. The tree is simulated by the Beta-Splitting model (see below), which allows producing imbalanced trees, consistent with what was observed in the real data [23].

On each edge (except for those attached to the root of the tree; see below), we simulate a number of CNAs, the number of which corresponds to a Poisson distribution (by default, *λ* = 2). *λ* relates with the mutation rate which has been studied for two decades [47, 66]. There has not yet been a comprehensive knowledge of the mutation rate of CNAs, but according to the data from [32], we found there are about several dozens of CNAs in this dataset. The same can be found in a pan-cancer study [82]. Setting default *λ* to be 2 will lead to the similar number of CNAs at the leaves for a tree. The daughter cell of the edge inherits all CNAs in the parent node, in addition to its unique CNAs. To simulate a CNA, we randomly choose the allele, and the chromosome and position on the allele that CNA is going to occur. First, we sample the allele on which the CNA is going to occur from the paternal and maternal alleles according to a binomial distribution (default *p* = 0.5). We designed the simulator in a framework which keeps track of the allele at which the CNA occurs so that in our future work of simulating single nucleotide variations (SNV) simultaneously, the allele that is dropped due to the high allelic dropout rate can be traced. For the CNA size, we sample from an exponential distribution (default mean=5Mbp), plus a minimum CNA size (default 2Mbp). We set the minimum CNA size by default to be 2Mbp because these CNAs are rare and commonly associated with disease [25], and also because of the limited resolution of single-cell data. The exponential distribution with mean 5Mbp is to render a wide range of CNA size. According to [34, 35], the larger the CNA size, the smaller the CNAs possibility. We choose copy number gain versus loss by a binomial distribution (default *p* = 0.5). We set the default parameter to be 0.5 so that copy number gain and loss are equally distributed. If a copy number gain is sampled, we sample from a geometric distribution (default *p* = 0.5) to determine the number of copies to be gained (mean=1*/p*). This choice of a distribution is motivated by the observation that extremely high copy number gains are very rare and are often observed by double minutes amplification [16], which we do not take into account currently. Once a whole-genome DNA sequence is simulated with the CNAs, the gained copies are placed in tandem with the original copy. If a copy number loss is sampled, the whole sequence on that region of the allele is deleted.

The CNAs on the edges attached to the root node are simulated differently. In particular, clonal whole chromosomal amplifications can occur on these edge, as indicated in the punctuated evolution model observed in [23]. We simulate the chromosomal amplifications in addition to the focal CNAs. We set the probability of a chromosome to be amplified to be according to a binomial distribution (default *p* = 0.2). This default value is used so that while the whole chromosome amplification is introduced, 20% chromosomes in the genome will be changed. The number of the amplified copy is sampled from a geometric distribution (default mean is *p* = 1) multiplied by a value (default is 1) to amplify the copy numbers simulated without changing the distribution. The distribution of the whole chromosomal amplification can be turned off as an option.

At the edge to the root, we also add an option to allow more CNAs than the other edges. This is again to mimic a scenario of punctuated evolution [23]. To do that, we sample a value from a Poisson distribution (by default, *λ* = 4) which is the multiplier of the average number of the CNAs that occur to the edges other than the root. Thus the edge to the root has on average 4 times (default parameter) more focal CNAs than those of other edges. The higher this number, the more focal CNAs the edge to the root carries. This parameter is introduced to allow the user to simulate data that mimics the punctuated evolution model. However, due to the diversity of models that have been summarized for cancer evolution [18], users can turn off this option or tune the parameter *λ* so that the simulated data corresponds to their observation and experience. In our study, the value of *λ* was chosen according to the length of the trunk observed in Fig. 6 in [23].

Once we have the tree and the DNA sequences for all leaf nodes, we simulate the generation of read data from the genomes. Given the coverage of the genome (by default 0.04X), the simulator divides the genome into non-overlapping bins each of which has a default size of 200,000bp. To simulate the coverage variability observed in single-cell data, we use a Markov Chain Monte Carlo (MCMC) Metropolis-Hastings algorithm to determine a sequence of numbers of read pairs to be sampled for each bin.

An input of the variability information is a point on the Lorenz curve, whose X axis represents the percentage of the reads, and Y axis represents the percentage of the coverage. We transform it to a Beta distribution by Equations (1) and (2) in [58]. Through this transformation, we can sample read counts from a Beta distribution that corresponds to the given Lorenz curve. The followings are mathematical equations in [58] that are used to calculate the parameters (*α* and *β*) for the Beta distribution. In more detail, suppose *X* is a random variable whose cumulative distribution function *F* corresponds to a Beta distribution with parameters *α* and *β*. A point *x* sampled from this distribution has its corresponding X and Y positions on the Lorenz curve as *F* (*x*) and *ϕ*(*x*), where

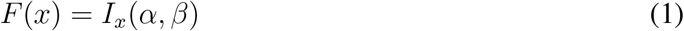

and

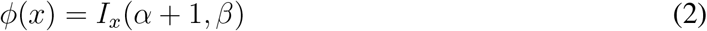

Given a point (*F* (*x*)*, ϕ*(*x*)) on the Lorenz curve, we can calculate *α* and *β* for the Beta distribution. Given the Beta distribution’s parameters, we can then sample read count for each bin by MCMC Metropolis-Hastings algorithm. Starting from the first bin whose read count is assigned as the expected coverage *x*_0_, we sample the next bin’s proposed read count *x^’^* by a Gaussian distribution, and accept it if compared with the previous bin’s read count *x*_0_,

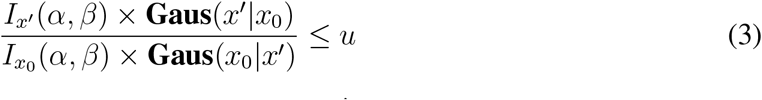

where **Gaus**(*x’|x*_0_) is the proposal probability of proposing *x’* given *x*_0_, and *u* is the acceptance ratio. We set *u* to be 0.5 by default. We set the same standard deviations for **Gaus**(*x’|x*_0_) and **Gaus**(*x*_0_*|x’*), centered at *x*_0_ and *x’*, respectively. Thus the two Gaussian distribution canceled out. The rest term, *I_x_’ /I_x_*_0_ ≤ *u*, controls how much the next bin’s read count differs from the current one. The read counts drawn are thus corresponding to a Beta distribution, and are simultaneously constrained by the acceptance ratio of the Metropolis-Hastings algorithm. This is to mimic the realistic data whose read coverage fluctuates, but the read count changes smoothly without sharp changes between neighboring bins.

### Running the programs

In all experiments, we eliminated reads that have mapping quality score *<* 40. We eliminated the cells that HMMcopy predicted as normal cells (predicted to be diploid and found no copy number aberration) in all experiments, the percentage of which was very small (*<* 0.2%).

### Parameters of simulator

The simulator is designed to be flexible, with user-specified parameters, as now describe.

#### Parameters for varying the ploidy level

To generate data with different ploidies, parameters associated with whole chromosomal amplification can be set for that purpose.

- **-W (–whole-amp)** Controls whether there are whole chromosomal amplifications or not.
- **-d (–del-rate)** The rate of copy number loss versus copy number gain.
- **-C (–whole-amp-rate)** The possibility that a chromosome is selected to have whole chromosomal amplification.
- **-E (–whole-amp-num)** For those chromosomes that are selected to be amplified, multiplying this number with the sampled value from a geometric distribution, whose *p* is “-J” described below, renders the final number of copies to be amplified.
- **-J (–amp-num-geo-par)** The parameter *p* in the geometric distribution from which the number of copy of the chromosome to be amplified is sampled. Combination of **-J** and **-E** can make a variety of copy number distributions and make it convenient to attain higher copy number gains when necessary.

In our experiment above where we varied the ploidy level, we use a combination of these five parameters to generate data whose ploidies range from 1.5 to 5 as follows.

- **Ploidy = 1.5**: **-W 0 -d 1** No amplifications are allowed, and all copy number aberrations come from deletion.
- **Ploidy = 3**: **-W 1 -d 0.5 -C 0.5 -E 1 -J 1** Amplification is allowed, and the average number of amplification for the whole genome is 0.5 for one allele. The final ploidy is 3.
- **Ploidy = 4**, the case that lacks odd copy numbers: **-W 1 -d 0.5 -C 0.5 -E 2 -J 1** Amplification is allowed, and the average number of amplification for the whole genome is 1 for one allele. The final ploidy is 4. Note that since the parameter *p* in the geometric distribution (-J) is set to be one, the copy number is amplified by two for the allele that is selected for amplification. This causes the lack of intermediate copy numbers such as three, five, etc.
- **Ploidy = 3.8**, the case that has odd copy numbers: **-W 1 -d 0.5 -C 0.9 -E 1 -J 1** Amplification is allowed, and the average number of amplification for the whole genome is 0.9 for one allele. The final ploidy is 3.8. Compared with the previous case which lacks odd copy numbers, we increase the copy number by doubling 90% of the chromosomes. The following local copy number aberrations that are performed based on the amplified genome will then generate regions that have different copies, including the odd copies. In the absence of odd copy numbers, copy numbers 2, 4 and 6 will be considered as 1, 2, and 3 by any method. Thus, without copy numbers 1, 3 and 5, there is no way for a method to tell the correct absolute copy number.
- **Ploidy = 5**, the case that lacks intermediate copy numbers: **-W 1 -d 0.5 -C 0.5 -E 3 -J 1** Amplification is allowed, and the average number of amplifications for the whole genome is 1 for one allele. The final ploidy is 5. Note that since the parameter *p* in the geometric distribution (-J) is set to be one, the copy number is amplified by three for the allele that is selected for amplification. This causes a scenario where most of the copy numbers are two, five and eight.
- **Ploidy = 5.26**, the case that has intermediate copy numbers: **-W 1 -d 0.5 -C 0.9 -E 1 -J 0.55**
- Amplification is allowed, and the average number of amplification for the whole genome is for one allele. Setting parameter *J* to be 0.55, the total amplified copy number for each allele is 1.63 (from 1*/p ×* 0.9). The final ploidy is 5.26.

#### Parameters for varying the read count distribution

Since Lorenz curves have been used to evaluate the variability of read counts [36, 75, 76], we used the Lorenz curves reported in [76] for simulating variabilities at different levels. We sampled the read counts for each bin by the distribution (Beta distribution) corresponding to their Lorenz curves using a Markov Chain Metropolis-Hastings method (Additional file 1: Figure S19 shows the Lorenz curves (left panel) and their corresponding Beta distributions (right panel) for the four technologies. The key parameters used for the Lorenz curves and Beta distributions corresponding to the four technologies are shown within the panels.).

### The Beta-splitting model

For generating the underlying evolutionary trees, we followed a generalization of the Blum-François Beta-splitting model [10] which is inspired by Aldous’ Beta-splitting model [2]. The construction of a tree based on this model [59] consists of two major steps: First, we generate two sequences of random values: *B* = (*b*_1_*, b*_2_*, · · ·*) and *U* = (*u*_1_*, u*_2_*, · · ·*), *B* is a sequence of independent and identically distributed (i.i.d.) random variables sampled from the *ℬ* (*α* + 1*, β* + 1) distribution, and, *U* is a sequence of i.i.d. random variables with the uniform distribution on [0, 1]. We call *g_i_* = (*u_i_, b_i_*)*_i∈_*_ℕ_ the *generating sequence* which is the basis of incremental construction of a tree. At the second step, we run the following algorithm on the random values generated at the first step. The process of constructing an evolutionary tree for *n* cells/leaves based on the Beta-splitting model with the parameters (*α, β*) combining these two steps is described in the following pseudocode.

#### Algorithm 1 Algorithm for constructing a tree *𝒥* with *n* leaves with the Beta parameters *α* and *β*.

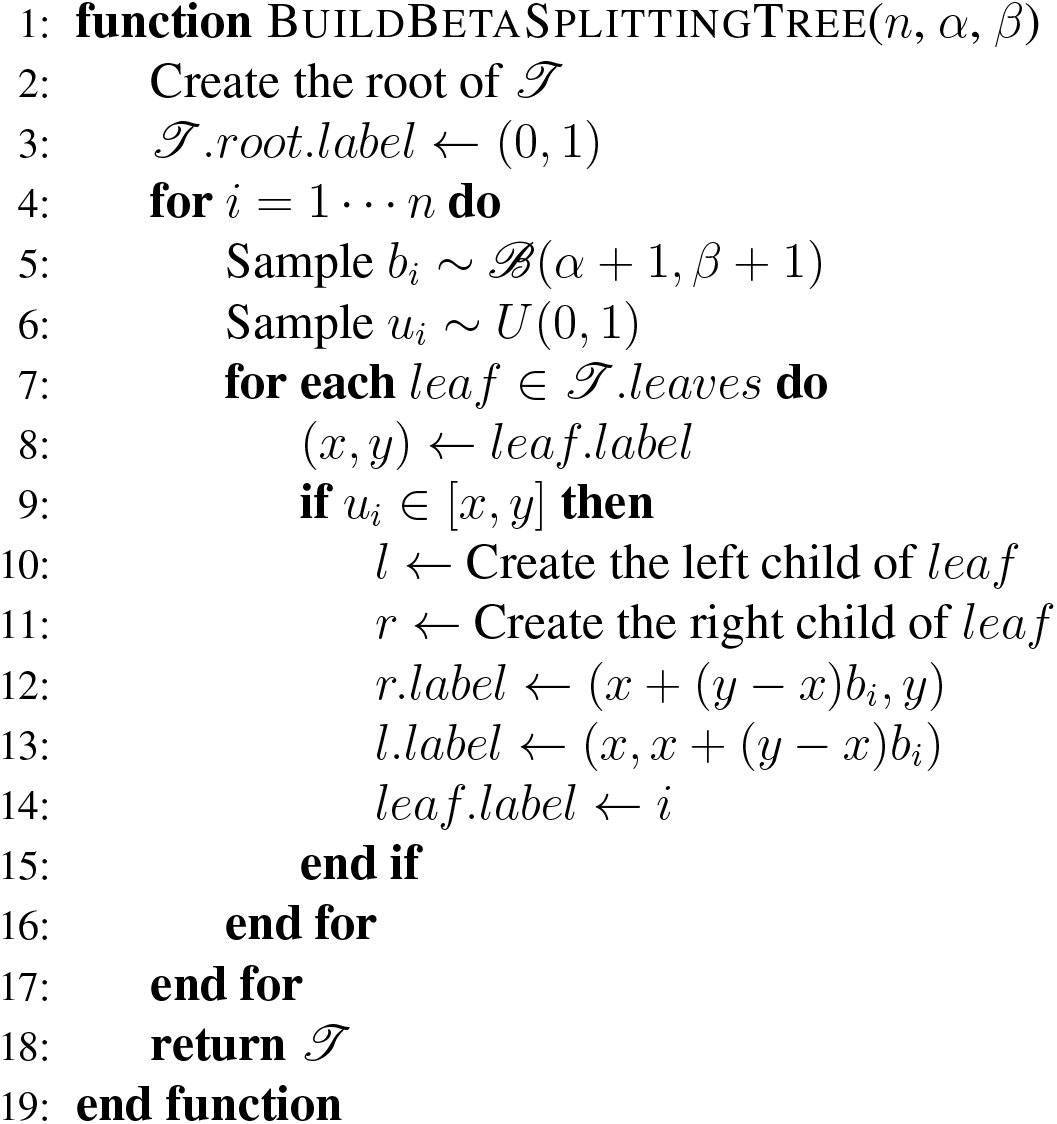

### Software availability

The simulator has been implemented in Python and is freely available at https://bitbucket.org/ xianfan/cnsc simulator/src/master/, which also includes the scripts for regenerating the comparison results for both simulated and real datasets.

## Availability of data and materials

The new version of HMMcopy was downloaded from https://github.com/shahcompbio/single cell pipeline/tree/master/single cell/workflows/hmmcopy. The scripts to preprocess files for HMM-copy were downloaded from https://shahlab.ca/projects/hmmcopy utils/. We use hg19 for all experiments in this manuscript and the mappability file used by HMMcopy was downloaded from http://genome.ucsc.edu/cgi-bin/hgFileUi?db=hg19&g=wgEncodeMapability. CopyNumber was downloaded from https://bioconductor.org/packages/release/bioc/html/copynumber.html. Ginkgo’s command line version which was used in this manuscript was downloaded from https://github.com/ robertaboukhalil/ginkgo.

The real biological dataset that we analyzed is available from NCBI Sequence Read Archive under accession SRP114962.

## Glossary

- ***Segmentation*** Computationally segmenting the genome into non-overlapping regions so that each region has a homogeneous copy number.
- ***Boundary*** and ***Breakpoint*** Positions on the genome where segmentation occurs.
- ***Absolute Copy Number*** The integer value representing the number of copies of a region on the genome.
- ***Ploidy*** The average copy number across the genome.

## Acknowledgement

The authors acknowledge the Texas Advanced Computing Center (TACC) at The University of Texas at Austin for providing HPC resources that have contributed to the research results reported within this paper. We thank Dr. Ruli Gao for her help in explaining the real dataset, and Mr. Alexander Davis for his insights on the ploidy of the single cells.

## Competing interests

The authors declare that they have no competing interests.

## Author’s contributions

XF, ME, NN, and LN designed the study. XF and ME wrote the code and ran the experiments. All authors wrote and approved the manuscript.

## Funding

The study was supported by the National Science Foundation grant IIS-1812822 (L.N.). X.F. was supported in part by a Computational Cancer Biology Training Program (CPRIT Grant No. RP170593).

## Notes

#### Summary of Updates

We have extended the manuscript to give a more comprehensive listing of the methods out there. However, it is important to note that a major part of this manuscript is still about benchmarking the three most commonly used methods (so it's not a "review" paper in that sense).

