## Supplementary Material for "Methods for Copy Number Aberration Detection from Single-cell DNA Sequencing Data" (from another version of this preprint: version 1, 2019-07-09)

### Supplementary material for
### Benchmarking Tools for Copy Number Aberration Detection from Single-cell DNA Sequencing Data

Xian Fan[1,†], Mohammadamin Edrisi[1,†], Nicholas Navin[2], Luay Nakhleh[1,*]

[1]Department of Computer Science, Rice University, Houston, Texas, USA

[2]Department of Genetics, the University of Texas M.D. Anderson Cancer Center, Houston, Texas 77030, USA

[†]Contributed equally

July 4, 2019

### 1 Supplemental Figures

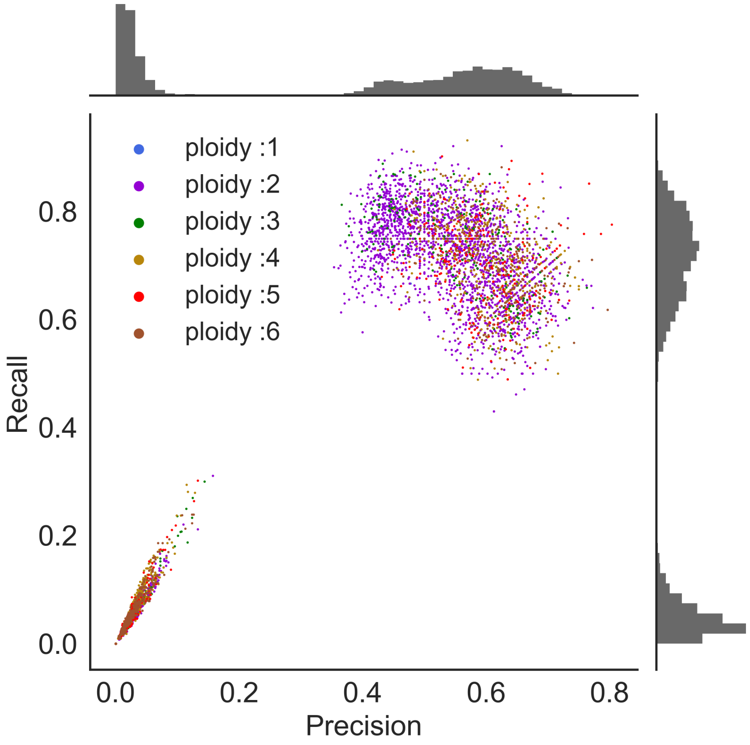


Figure S1: Scatter plot of precision and recall of HMMcopy on the simulated large dataset. The color of the dots represent the ploidy that HMMcopy predicts. The upper and right histograms show the histogram of precision and recall, respectively, for all cells.

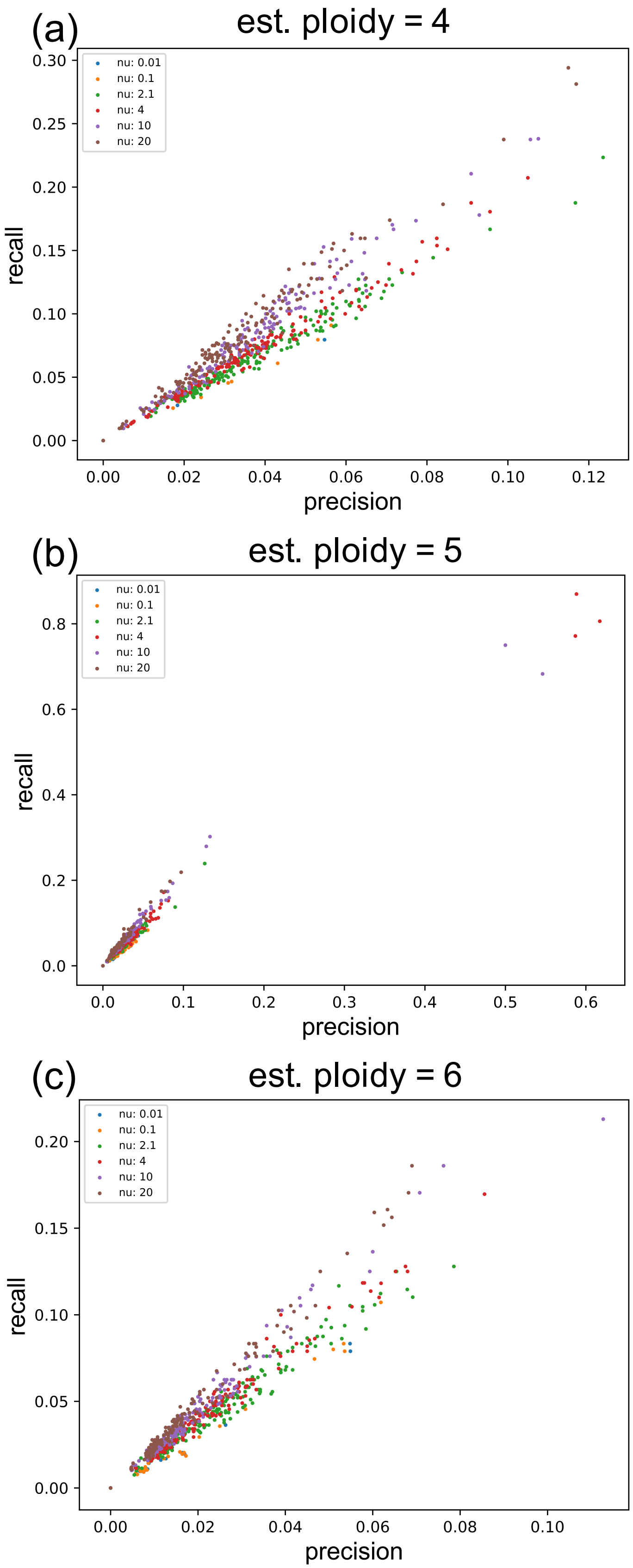


Figure S2: Precision and recall of HMMcopy on the simulated large dataset when ploidy estimation deviates from the ground truth which is 2.6: (a), (b) and (c) correspond to estimated ploidy 4, 5 and 6, respectively.

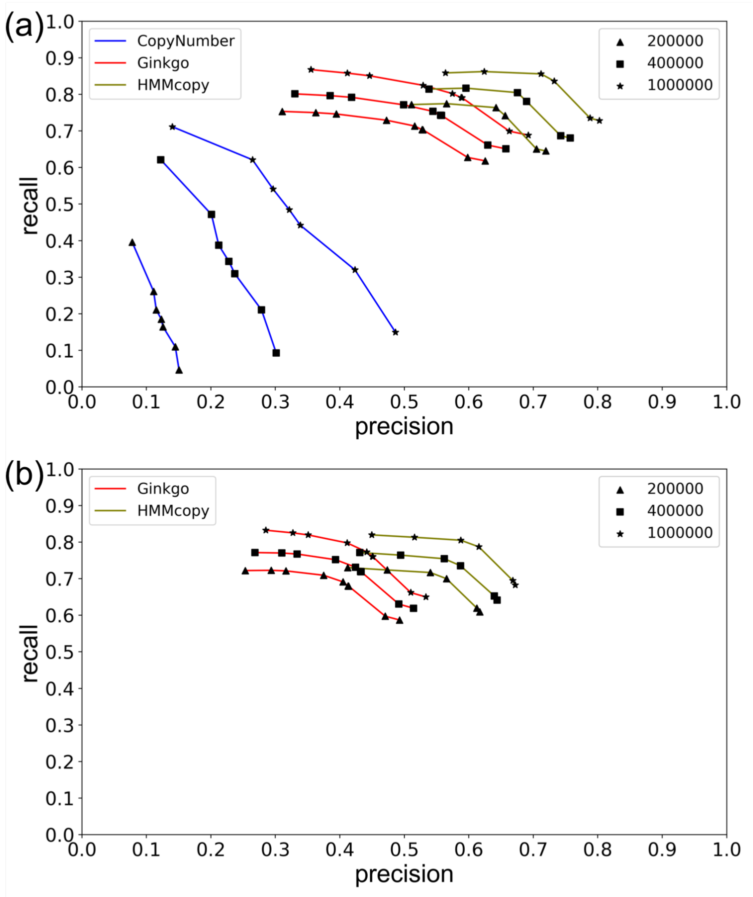


Figure S3: ROC curves of the three methods HMMcopy, Ginkgo, and CopyNumber on a second repetition. (a) coarse-grained analysis results, and (b) fine-grained results for the second repetition. For each method, the results based on three thresholds of correctness are plotted. For HMMcopy, $nu$, which controls the suggested degree of freedom between states, was tuned to take on the values 0.01 (rightmost), 0.1, 2.1 (the tool's default), 4, 10, and 20 (leftmost). For Ginkgo, $alpha$, which controls the significance level to accept a change point, was tuned to take on the values 1e-1000 (rightmost), 1e-100, 1e-10, 1e-5, 1e-4, 1e-3, 1e-2 (the tool's default), 0.02, and 0.05 (leftmost). The dots corresponding to values 1e-5 and 1e-10 in coarse-grained analysis overlap. For CopyNumber, $gamma$, which is the weight of the penalty on changing a state, was tuned to take on the values 40 (rightmost, and the tool's default), 10, 5, 4, 3, 2, and 1 (leftmost).

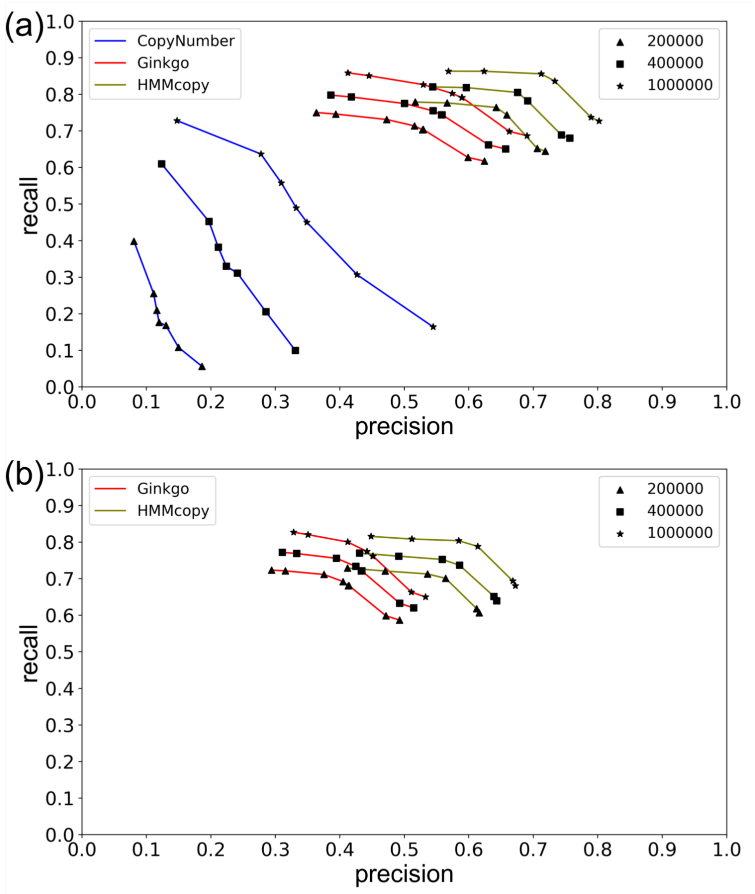


Figure S4: ROC curves of the three methods HMMcopy, Ginkgo, and CopyNumber on a third repetition. (a) Coarse-grained analysis results, and (b) fine-grained results for the third repetition. For each method, the results based on three thresholds of correctness are plotted. For HMMcopy, $nu$, which controls the suggested degree of freedom between states, was tuned to take on the values 0.01 (rightmost), 0.1, 2.1 (the tool's default), 4, 10, and 20 (leftmost). For Ginkgo, $alpha$, which controls the significance level to accept a change point, was tuned to take on the values 1e-1000 (rightmost), 1e-100, 1e-10, 1e-5, 1e-4, 1e-3, 1e-2 (the tool's default) and 0.02 (leftmost). The dots corresponding to values 1e-5 and 1e-10 in coarse-grained analysis overlap. For CopyNumber, $gamma$, which is the weight of the penalty on changing a state, was tuned to take on the values 40 (rightmost, and the tool's default), 10, 5, 4, 3, 2, and 1 (leftmost).

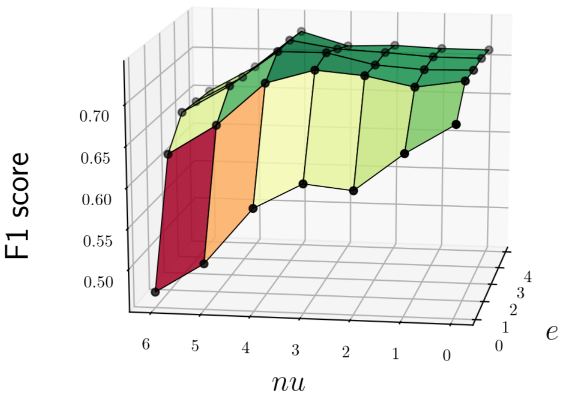


Figure S5: Plot of F1 scores of HMMcopy based on varying *nu* and *e* on one simulated dataset (1000 cells). On *nu*, 0, 1, 2, 3, 4, 5 and 6 represent 0.01, 0.1, 1, 2.1 (default), 4, 10 and 20. On *e*, 0, 1, 2, 3, and 4 represent 0.99, 0.9999, 0.999999, 0.99999999 (default), and 0.9999999999. It can be seen when *nu* is 4, and *e* is $\geq$ 0.999999, F1 score reaches the maximum.

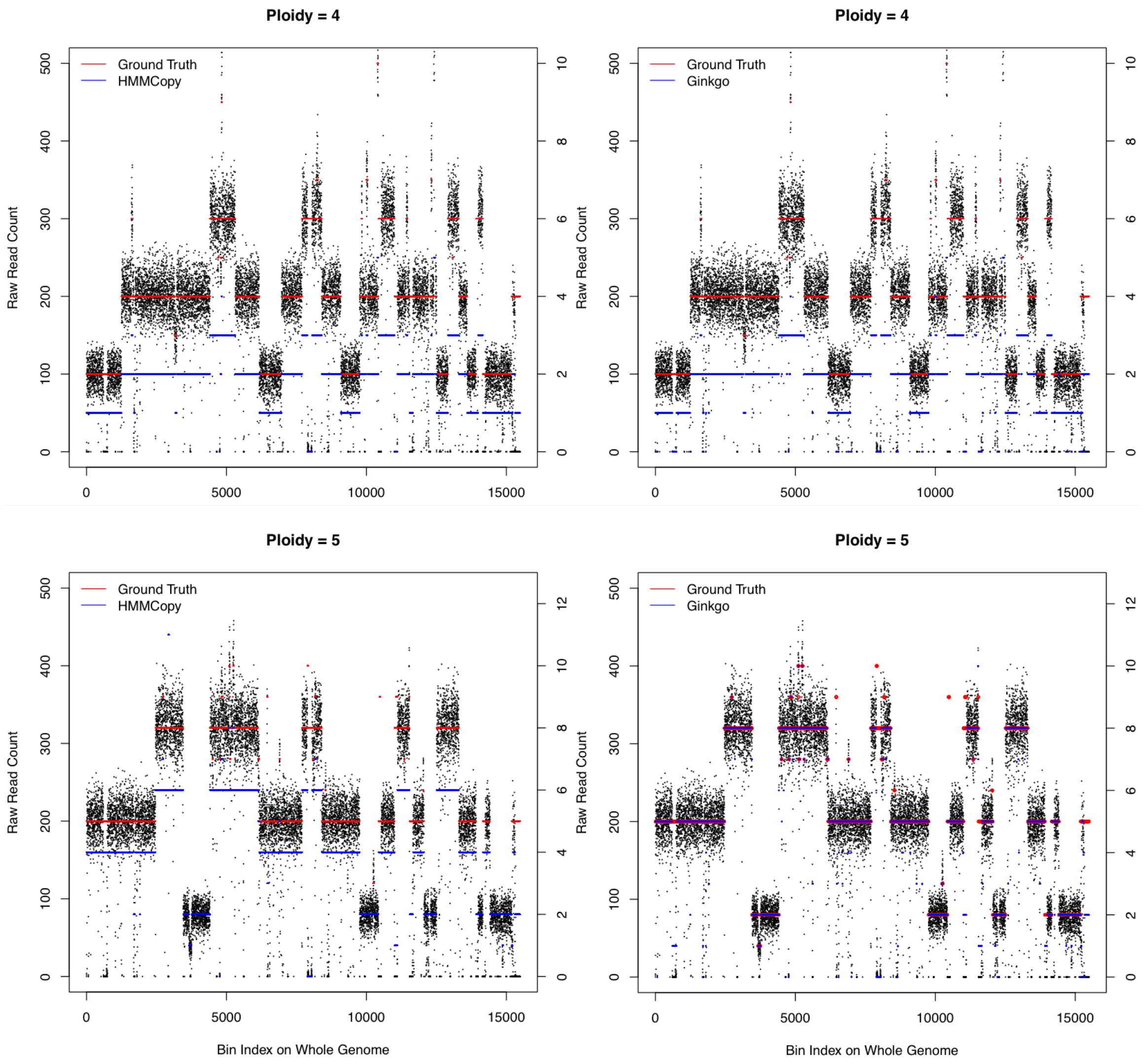


Figure S6: Example of two cells that lack odd copy numbers (top panels, simulated ploidy=4) and intermediate copy numbers (bottom panels, simulated ploidy=5), respectively. The left and right panels show HMMcopy and Ginkgo's results, respectively. For each panel, X and primary Y axes represent the whole genome segmented into bins and the raw read count for each bin, respectively. The secondary Y axis represents the absolute copy number. Black dots represent the raw read count for each bin. Red and blue horizontal lines represent the ground truth of the absolute copy number and the inferred ones from HMMcopy and Ginkgo. Due to the lack of odd copy numbers, both HMMcopy and Ginkgo incorrectly predicted absolute copy number for the case of ploidy=4. Due to the lack of intermediate copy number, HMMcopy incorreclty predicted absolute copy number for the case of ploidy=5. Despite the lack of intermediate copy numbers, Ginkgo correctly predicted absolute copy number for the case of ploidy=5 (red and blue lines overlap in this case).

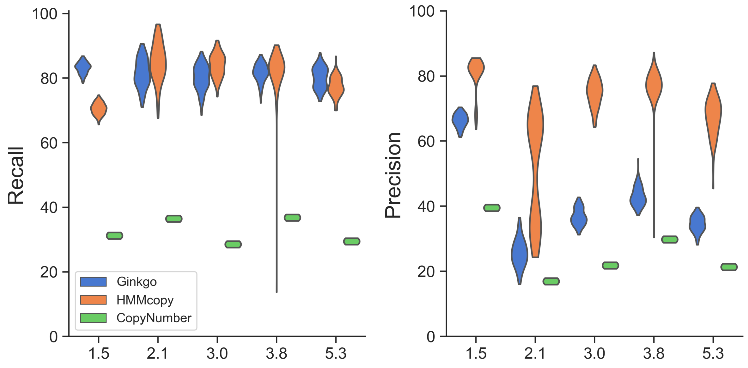


Figure S7: Recall and Precision of Ginkgo, HMMcopy, and CopyNumber on varying ploidy level based on the coarse-grained analysis on a second repetition. The ploidies of the simulated data were 1.5, 2.1, 3.0, 3.8, and 5.3.

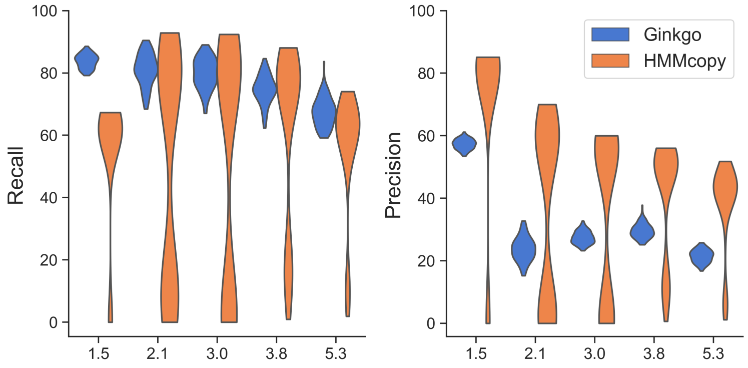


Figure S8: Recall and Precision of Ginkgo and HMMcopy on varying ploidy level based on the fine-grained analysis on a second repetition. The ploidies of the simulated data were 1.5, 2.1, 3.0, 3.8, and 5.3.

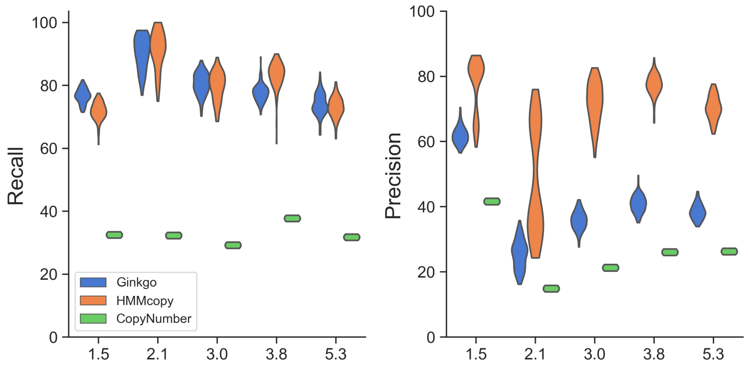


Figure S9: Recall and Precision of Ginkgo, HMMcopy, and CopyNumber on varying ploidy level based on the coarse-grained analysis on a third repetition. The ploidies of the simulated data were 1.5, 2.1, 3.0, 3.8, and 5.3.

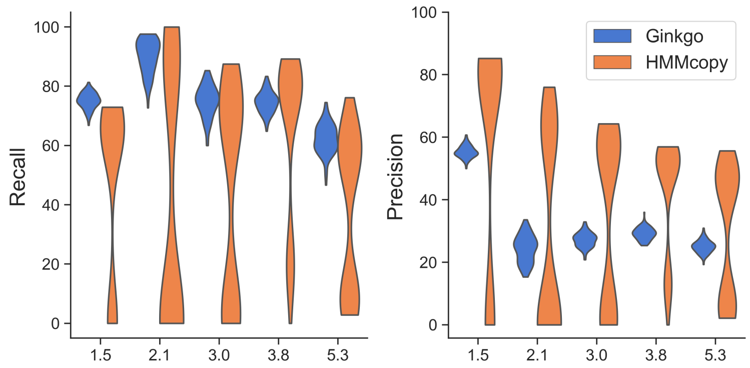


Figure S10: Recall and Precision of Ginkgo and HMMcopy on varying ploidy level based on the fine-grained analysis on a third repetition. The ploidies of the simulated data were 1.5, 2.1, 3.0, 3.8, and 5.3.

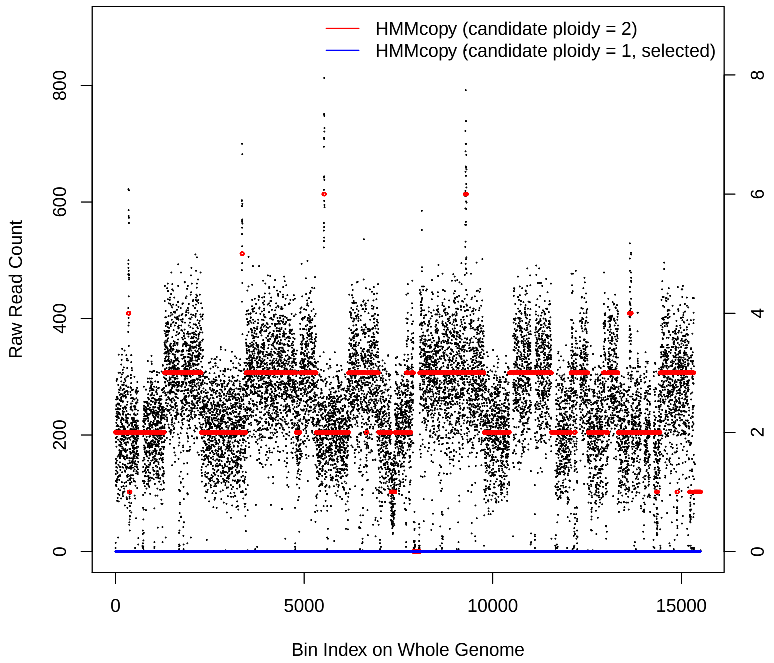


Figure S11: An example of when wrong selection of ploidy in HMMcopy leads to wrong estimation of breakpoints. X and primary Y axes represent the whole genome segmented into bins and the raw read count for each bin, respectively. The secondary Y axis represents the absolute copy number. Black dots represent the raw read count for each bin. Red and blue horizontal lines represent the copy number profiles of HMMcopy on the condition when its ploidy estimation is 2 and 1, respectively, the latter of which was selected as the optimal ploidy by HMMcopy.

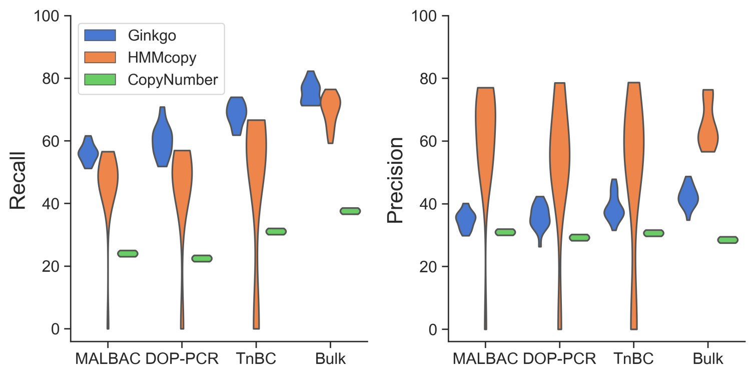


Figure S12: Recall and Precision of Ginkgo, HMMcopy, and CopyNumber on varying coverages based on the coarse-grained analysis on a second repetition. The coverages are varied to mimic those produced by MALBAC, DOP-PCR, TnBC and Bulk sequencing.

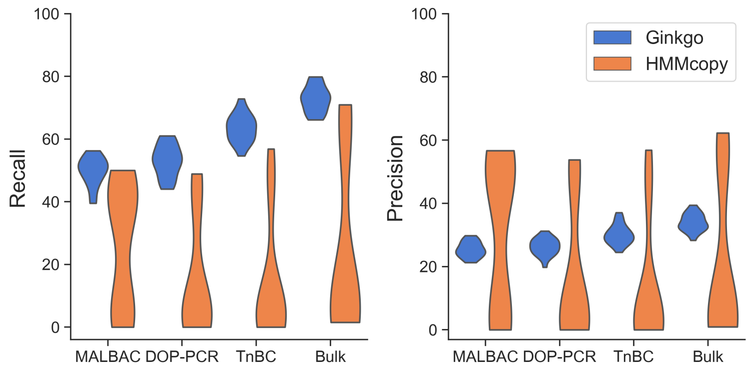


Figure S13: Recall and Precision of Ginkgo and HMMcopy on varying coverages based on the fine-grained analysis on a second repetition. The coverages are varied to mimic those produced by MALBAC, DOP-PCR, TnBC and Bulk sequencing.

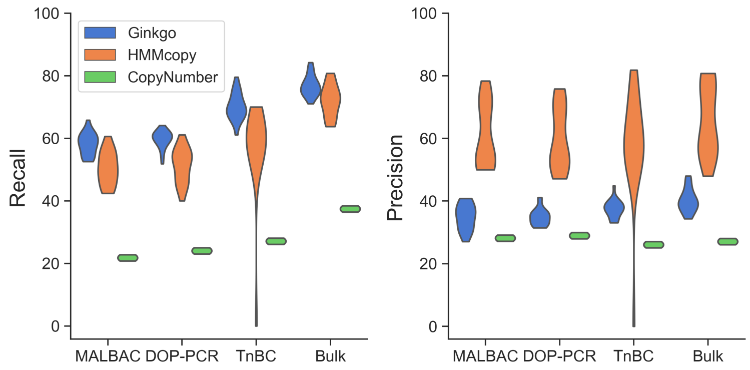


Figure S14: Recall and Precision of Ginkgo, HMMcopy, and CopyNumber on varying coverages based on the coarse-grained analysis on a third repetition. The coverages are varied to mimic those produced by MALBAC, DOP-PCR, TnBC and Bulk sequencing.

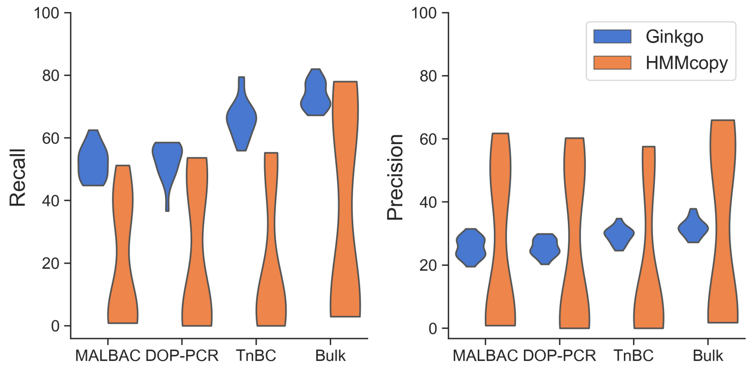


Figure S15: Recall and Precision of Ginkgo and HMMcopy on varying coverages based on the fine-grained analysis on a third repetition. The coverages are varied to mimic those produced by MALBAC, DOP-PCR, TnBC and Bulk sequencing.

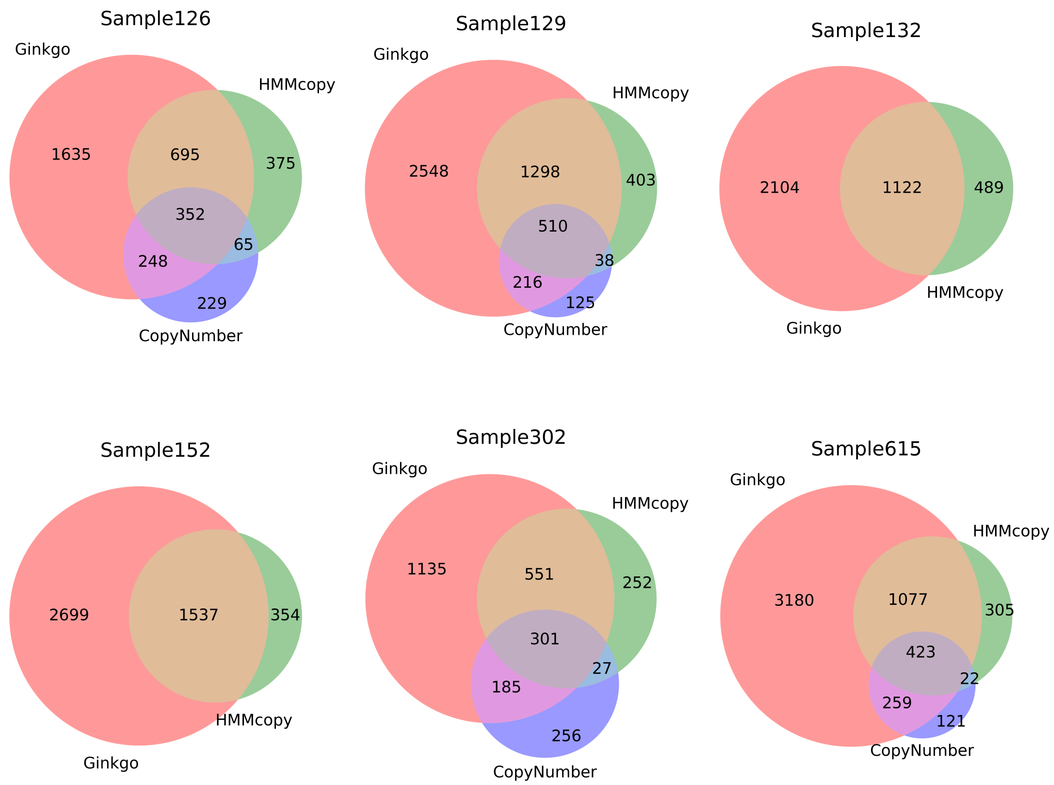


Figure S16: Venn diagram of the breakpoints from Ginkgo, HMMcopy and CopyNumber for the rest of the six samples. Breakpoints from two methods are counted as overlapped ones if they are within 400,000bp.

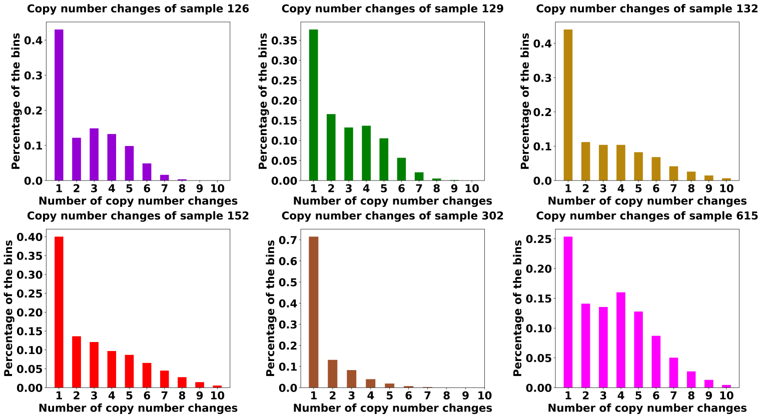


Figure S17: Distribution of the copy number changes (under a parsimony analysis) per bin based on the copy number profiles obtained by HMMcopy for the seven samples. A maximum parsimony tree was inferred from the copy number profiles of the cells, and the minimum number of copy number changes per bin along all branches of the tree was computed by parsimony analysis. The percentages of bins with each number of copy number changes are plotted.

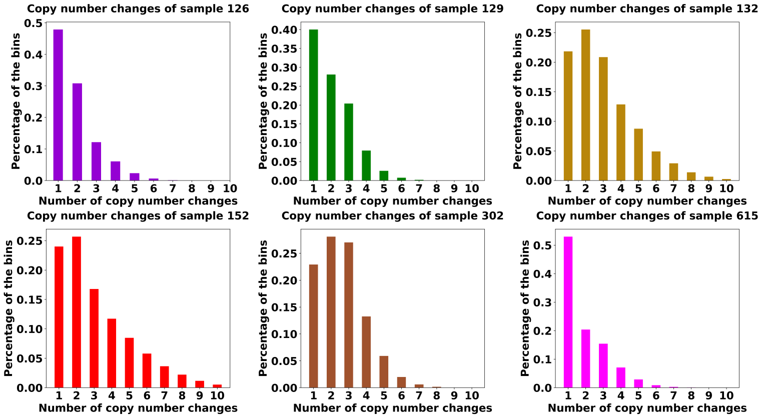


Figure S18: Distribution of the copy number changes (under a parsimony analysis) per bin based on the copy number profiles obtained by Ginkgo for the seven samples. A maximum parsimony tree was inferred from the copy number profiles of the cells, and the minimum number of copy number changes per bin along all branches of the tree was computed by parsimony analysis. The percentages of bins with each number of copy number changes are plotted.

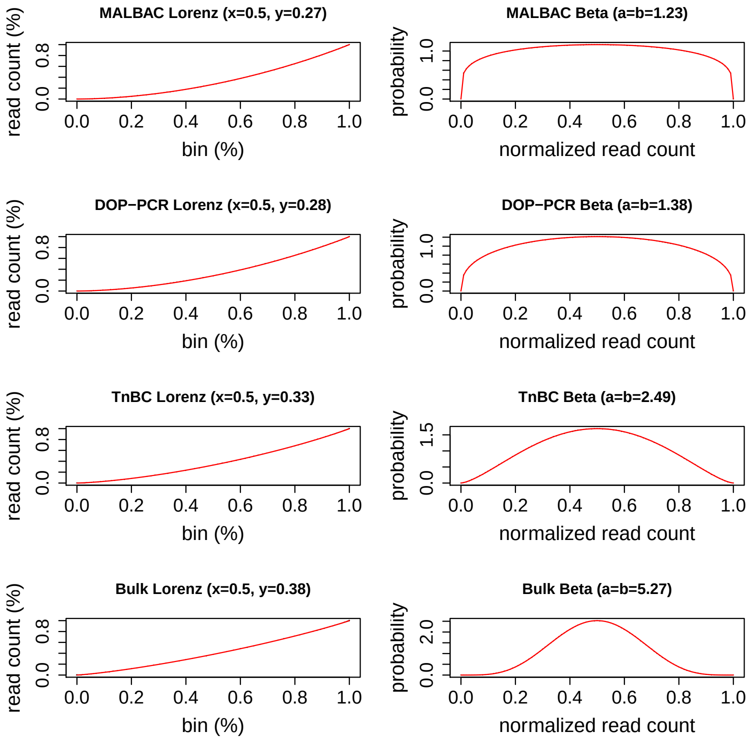


Figure S19: Left and right panels show the Lorenz curves and Beta distributions, respectively. The four rows represent four sequencing technologies: MALBAC, DOP-BAC, TnBC and Bulk. The parameters of generating Lorenz curve were learned from [1]. The parameters of generating Beta distributions were learned from Eq.1 and 2 in the main text.
